# Conformational dynamics of auto-inhibition in the ER calcium sensor STIM1

**DOI:** 10.1101/2020.12.17.423361

**Authors:** Stijn van Dorp, Ruoyi Qiu, Ucheor B. Choi, Minnie M. Wu, Michelle Yen, Michael Kirmiz, Axel T. Brunger, Richard S. Lewis

## Abstract

The dimeric ER Ca^2+^ sensor STIM1 controls store-operated Ca^2+^ entry (SOCE) through the regulated binding of its CRAC activation domain (CAD) to Orai channels in the plasma membrane. In resting cells, the STIM1 CC1 domain interacts with CAD to suppress SOCE, but the structural basis of this interaction is unclear. Using single-molecule Förster resonance energy transfer (smFRET) and protein crosslinking approaches, we show that CC1 interacts dynamically with CAD in a domain-swapped configuration with an orientation predicted to sequester its Orai- binding region adjacent to the ER membrane. Following ER Ca^2+^ depletion and release from CAD, cysteine crosslinking indicates that the two CC1 domains become closely paired along their entire length in the active Orai-bound state. These findings provide a structural basis for the dual roles of CC1: sequestering CAD to suppress SOCE in resting cells and propelling it towards the plasma membrane to activate Orai and SOCE after store depletion.

## Introduction

Store-operated Ca^2+^ entry (SOCE) is a nearly ubiquitous signaling pathway activated by extracellular stimuli that deplete Ca^2+^ from the endoplasmic reticulum (ER). SOCE is essential for many physiological functions of excitable and non-excitable cells, including gene expression, secretion, and motility (Prakriya and Lewis, 2015). Accordingly, loss-of-function mutations in the core SOCE components lead to serious human pathologies, including severe combined immunodeficiency, autoimmunity, myopathy, ectodermal dysplasia and anhidrosis (Lacruz and Feske, 2015), while gain-of-function mutations cause Stormorken syndrome, miosis, myopathy, thrombocytopenia and excessive bleeding (Böhm and Laporte, 2018). These clinical manifestations underscore the need for precise regulation of SOCE to ensure that it is silent when ER Ca^2+^ stores are full yet reliably activated by stimuli that drive store depletion.

The ER membrane protein STIM1 (Figure 1A, top) controls SOCE through the regulated interaction of its CRAC activation domain (CAD (Park et al., 2009); also known as SOAR (Yuan et al., 2009) or CCb9 (Kawasaki et al., 2009)) with Orai1 Ca^2+^ channels in the plasma membrane. In resting cells, Ca^2+^ bound to the luminal EF hands of the STIM1 dimer suppresses STIM1 activity by promoting intramolecular association of the coiled-coil 1 (CC1) domains with CAD, referred to as the ‘inhibitory clamp’ (Korzeniowski et al., 2010; Fahrner et al., 2014; Ma et al., 2015; Yang et al., 2012; Yu et al., 2013). Upon ER Ca^2+^ depletion, dissociation of Ca^2+^ from the luminal EF hands triggers conformational changes that release CC1 from CAD and promotes STIM1 accumulation at ER-plasma membrane (ER-PM) junctions where the CAD domain binds, traps, and activates Orai1 channels diffusing in the PM (Liou et al., 2007; Wu et al., 2014, 2006). Thus, to elucidate the regulatory mechanism of SOCE, it is essential to understand how CC1 interacts with CAD to control STIM1 activity.

**Figure 1.**
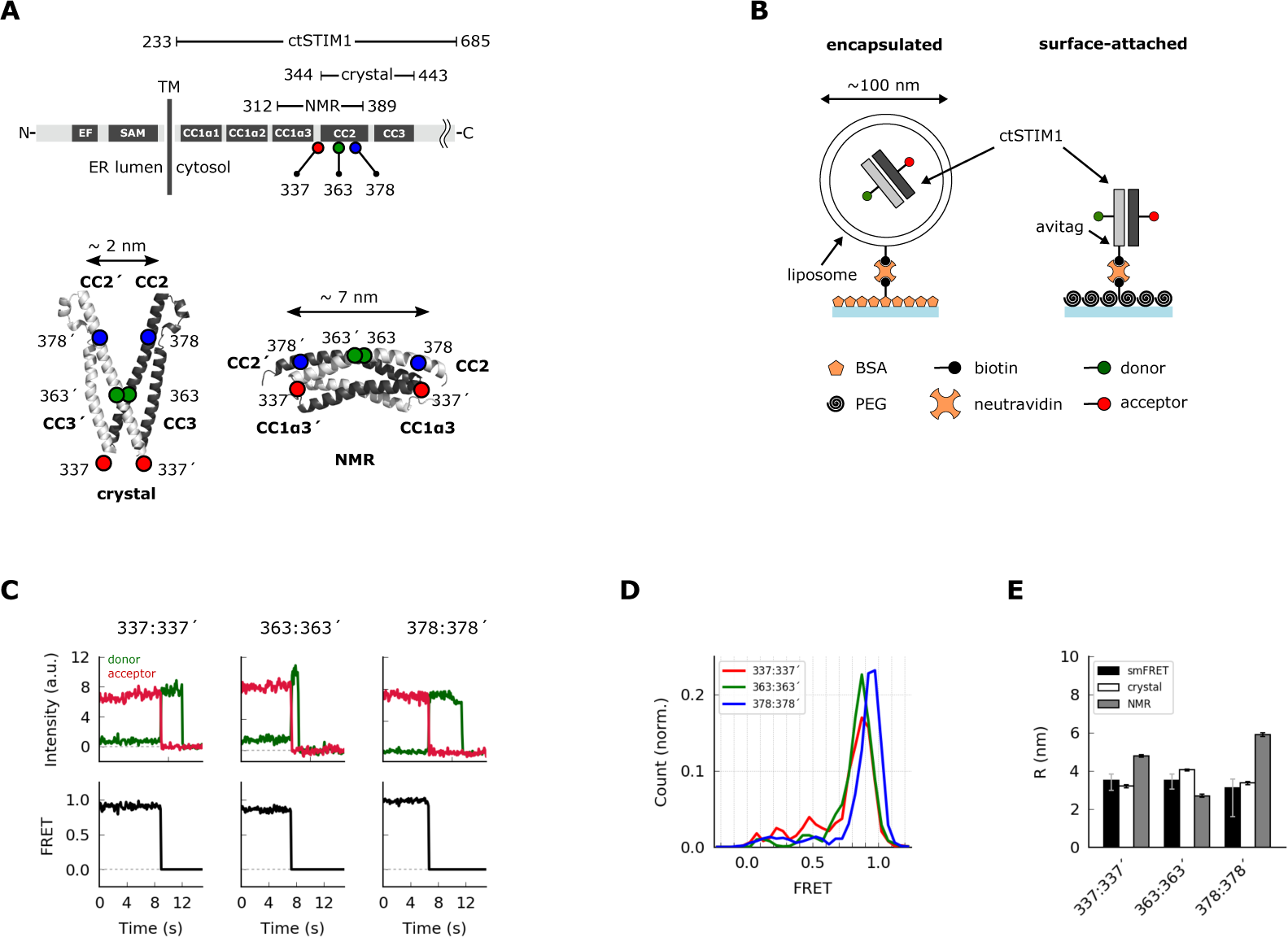
Parallel orientation of CC2 domains in ctSTIM1 is consistent with the CAD crystal structure. (**A**) (*top*) Schematic overview of domains in ctSTIM1, comprising the membrane-proximal coiled-coil 1 (CC1) domain (with helical regions α1, α2, and α3), and the CC2 and CC3 domains which are part of CAD. (*bottom*) The orientations of the CC2 domains are a prominent feature that distinguishes the crystal structure of CAD (*bottom left*, parallel CC2 domains; 3TEQ.pdb) and the NMR structure of CC1α3-CC2 (*bottom right*, anti-parallel CC2 domains; 2MAJ.pdb). Sites tested by inter-subunit smFRET are indicated. Although aa 337 is not part of the crystal structure (aa 344-443), it is sufficiently close to permit comparison to the NMR structure. (**B**) Two methods for immobilizing single ctSTIM1 dimers for smFRET. ctSTIM1 peptides were encapsulated in liposomes and immobilized on a glass slide by a biotin- neutravidin interaction or were attached directly via a C-terminal avitag. (**C**) Example recordings of donor (*green*) and acceptor (*red*) fluorescence evoked by donor excitation for sites 337:337’, 363:363’, and 378:378’. Single-step bleaching events indicate a single molecule, while the anti-correlated response of the donor to the acceptor bleach event is a hallmark of smFRET. Calculated instantaneous FRET ratios (*black*) are shown below. (**D**) Ensemble smFRET histograms for sites 337:337’, 363:363’, and 378:378’ displayed predominantly high FRET, indicating close parallel apposition. (**E**) A comparison of smFRET-derived distances (*black*) with simulated inter-fluorophore distances from the crystal (*white*) and NMR structures (*gray*). The measured distances closely match the crystal structure but deviate strongly from the NMR structure, particularly for 337:337’ and 378:378’. smFRET error bars indicate the expected distance deviation corresponding to an uncertainty of ±0.05 in FRET measurement. Simulation error bars indicate the s.e.m. of inter-fluorophore distance from 100 simulations of dye position (see Methods).

The CC1 domain comprises three helical segments, CC1α1-3 (Soboloff et al., 2012) (Figure 1A). Current evidence suggests that CC1α1, the most ER-proximal segment, acts as a bimodal switch to stabilize both the quiescent and active states of STIM1. When ER Ca^2+^ stores are full, CC1α1 binds to the CAD CC3 helix to sequester CAD (Fahrner et al., 2014; Ma et al., 2015); upon store depletion, the luminal EF-SAM domains dimerize, CC1α1 unbinds from CAD, and the transmembrane and proximal CC1α1 regions form a coiled-coil (Stathopulos et al., 2006; Schober et al., 2019; Hirve et al., 2018), projecting CAD towards the plasma membrane.

Although several residues in CC1α1 and CC3 have been identified as essential for forming the inhibitory clamp (Fahrner et al., 2014; Muik et al., 2011; Zhou et al., 2013; Ma et al., 2015), the helical arrangement of the CC1α1:CAD binding interface is completely unknown. In addition, while CC1α3 is implicated in stabilizing the inactive state (Yang et al., 2012; Korzeniowski et al., 2010), it is unclear whether this results from binding to CAD or another mechanism (Zhou et al., 2013).

Attempts to determine the structure of full-length STIM1 in the inactive or active state have been unsuccessful. Instead, structures of several fragments from the STIM1 cytosolic region have offered potential clues, although it is not yet clear how they might relate to physiological states of the full-length protein. The crystal structure of human STIM1 CAD (aa 345-444, with mutations L374M, V419A, and C437T) depicts a parallel V-shaped dimer comprising CC2 and CC3 domains (Figure 1A, bottom left) (Yang et al., 2012). It is unknown whether this structure represents an active or inactive conformation, and the absence of CC1 precludes inferences about its interaction with CAD. A solution NMR structure of a CC1α3- CC2 fragment depicts a dimer of wedged antiparallel CC2 helices (Figure 1A, bottom right) (Stathopulos et al., 2013). This NMR structure can bind Orai1 C-terminal fragments *in vitro* and has been proposed to be an intermediate in the activation of STIM1 (Stathopulos et al., 2013), but to our knowledge it has never been shown to bind intact Orai1 channels *in situ* and lacks the CC3 helix needed to evoke SOCE (Covington et al., 2010). Finally, in a crystal structure of CC1 (aa 237-340, with mutations M244L and L321M), two elongated CC1 helices form an antiparallel coiled-coil dimer within the CC1α2 and CC1α3 domains (see Figure 5D (Cui et al., 2013), but it is unclear how this elongated structure with its widely separated C-termini could attach to the closely paired N-termini of CAD.

**Figure 5.**
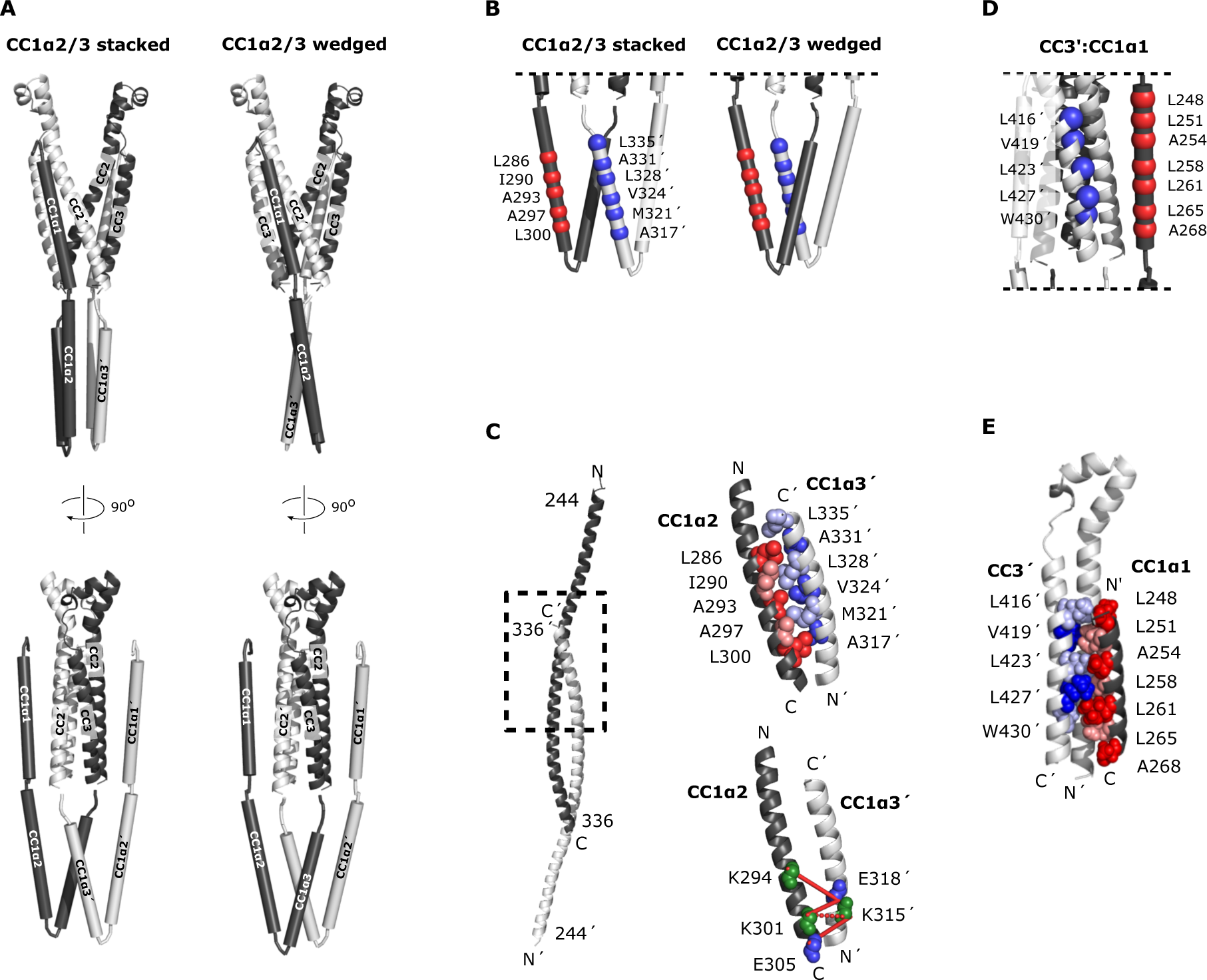
A model of the configuration of CC1 and CAD in ctSTIM1. (**A**) 36 smFRET- derived distances were used to reconstruct the orientations of CC1 domains relative to CAD, yielding two classes of solutions with the CC1α2/3 domains in a ’stacked’ (*left*) or ’wedged’ (*right*) configuration. In both classes, CC1α1 domains are in close parallel apposition to the CC3 domains on the adjacent subunit in a domain-swapped configuration, with the CC1α2 and CC1α3 domains forming a compact parallel structure directed away from CAD. The average of 50 model solutions is shown for each class (see Supplementary Figure 4B for individual solutions). (**B**) smFRET-derived models suggest hydrophobic stabilization of the CC1α2/3 complex by antiparallel apposition of CC1α2 and CC1α3’. (**C**) The crystal structure of CC1 peptides defines an antiparallel interaction of CC1α2/α3’ domains (*left*, *dashed box*), in which heptad-like hydrophobic sequences form a tightly packed dimer interface (*top right*, adapted from 4O9B.pdb). (*bottom right*) Tight antiparallel packing of CC1α2/α3’ in ctSTIM1 was confirmed by inter-subunit crosslinks with EDC (*solid lines*) and BS^3^ (*dashed line*) (see also Supplementary Figure 6). (**D**) Parallel apposition of hydrophobic residues on CC1α1 and CC3’. Several of these were previously identified by mutagenesis to stabilize the inactive state of STIM1, including L248, L251, L416, and L423. (**E**) A model of the CC1α1:CC3’ interface obtained by computational peptide docking (see Methods). In the lowest-energy solution, the helices engage in direct mutual hydrophobic interactions.

Although the sequences of these three structures partially overlap, their conformations are incompatible with each other, raising fundamental questions about their potential relation to the inactive and active forms of STIM1. What is the conformation of CAD in the context of the entire cytosolic domain? How is CC1 arranged relative to CAD in the inactive state, and how does this change during activation? Which regions of STIM1 are stable, and which are flexible? To address these questions, we examined the structure and dynamics of the full cytosolic domain of STIM1 (ctSTIM1, aa 233-685) using single-molecule Förster resonance energy transfer (smFRET) measurements at sites throughout CC1 and CAD. smFRET is commonly applied to estimate intramolecular distances and reveal asynchronous molecular transitions that are typically obscured in population measurements. Our results show that CAD in ctSTIM1 resembles the CAD crystal structure, but with unexpected structure and flexibility in an apical region that is critical for Orai binding and activation. CC1α1 interacts with CAD in a domain-swapped configuration to form the inhibitory clamp, enabled by the arrangement of the CC1α2 and CC1α3 domains. The Stormorken mutation R304W activates ctSTIM1 by releasing CC1α1 from CAD. In studies of full-length STIM1 in intact cells, state-dependent cysteine crosslinking after store depletion suggests that after unbinding from CAD, the two CC1 domains pack together along their entire length, in contrast to the CC1 crystal and CC1α3-CC2 NMR structures. Together, these findings offer the first structural view of the STIM1 inhibitory clamp and reveal the massive conformational changes evoked by store depletion, which serve to reorient and translocate CAD towards the PM where it can bind Orai1 and activate SOCE.

## Results

### The CC2 domains in ctSTIM1 are oriented in a parallel configuration

We initially applied smFRET to test whether the conformation of CAD within the cytosolic domain (ctSTIM1, aa 233-685) resembles that of the CC2-CC3 crystal structure (Yang et al., 2012) or the CC1α3-CC2 NMR structure (Stathopulos et al., 2013), referred to below as the ‘crystal structure’ or ‘NMR structure,’ respectively. A major distinction between the two structures is that the CC2 domains (aa 345-391) in CAD are parallel in the crystal structure but antiparallel in the NMR structure (Figure 1A). After removing the single native cysteine in ctSTIM1 with a C437S mutation, we created three mutants with cysteines at positions 337, 363, and 378, for which the two structures predicted very different inter-subunit distances, particularly for 337:337’ and 378:378’ (hereafter ‘:’ is used to denote a pair of sites in the dimer and ‘’’ indicates a site on the adjacent subunit) (Figure 1A). The ctSTIM1 mutants were expressed in *E. coli* and purified, yielding full-length protein of >90% purity (see Methods). After randomly labeling the cysteine-mutant ctSTIM1 molecules with FRET donor (Alexa Fluor 555) and acceptor (Alexa Fluor 647) dyes, single ctSTIM1 dimers were surface-tethered to a glass coverslip for imaging by TIRF microscopy, either encapsulated in liposomes or attached directly through a tag on the C-terminus (Figure 1B; the attachment method for each measurement is shown in Supplementary Figure 1). Upon excitation of the donor dye, the donor and acceptor emissions of individual molecules were ratioed to calculate FRET efficiencies, which were high and stable at all three label sites (Figure 1C, D). These results are consistent with a CAD conformation in which the CC2 domains are closely apposed and parallel, as in the crystal structure. Inter-fluorophore distances calculated from simulations of the donor and acceptor dye positions on the crystal structure (see Methods) closely matched those derived from smFRET, while simulations based on the NMR structure deviated substantially (Figure 1E). We conclude that in the context of the complete ctSTIM1 in solution, the CC2 domains in CAD adopt a parallel conformation consistent with the CAD crystal structure and not the NMR structure.

### The apex of CAD deviates from the crystal structure

We made additional smFRET measurements throughout the entire CAD to allow a more complete comparison with the crystal structure. Near the C-terminal end of the CC3 domain (aa 408-437), the 431:431’ label pair produced a stable, narrowly defined high-FRET state (Figure 2A), and inter-subunit smFRET throughout the CAD exceeded 0.5, consistent with a compact parallel dimer (Figure 2B). Near the dimer interface (hereafter termed the ‘base’ of CAD), smFRET-derived distances quantitatively agreed with those from the crystal structure, and an intra-subunit measurement 431:363 directly confirmed the compact folding of CC3 onto CC2 within each subunit (Figure 2C).

**Figure 2.**
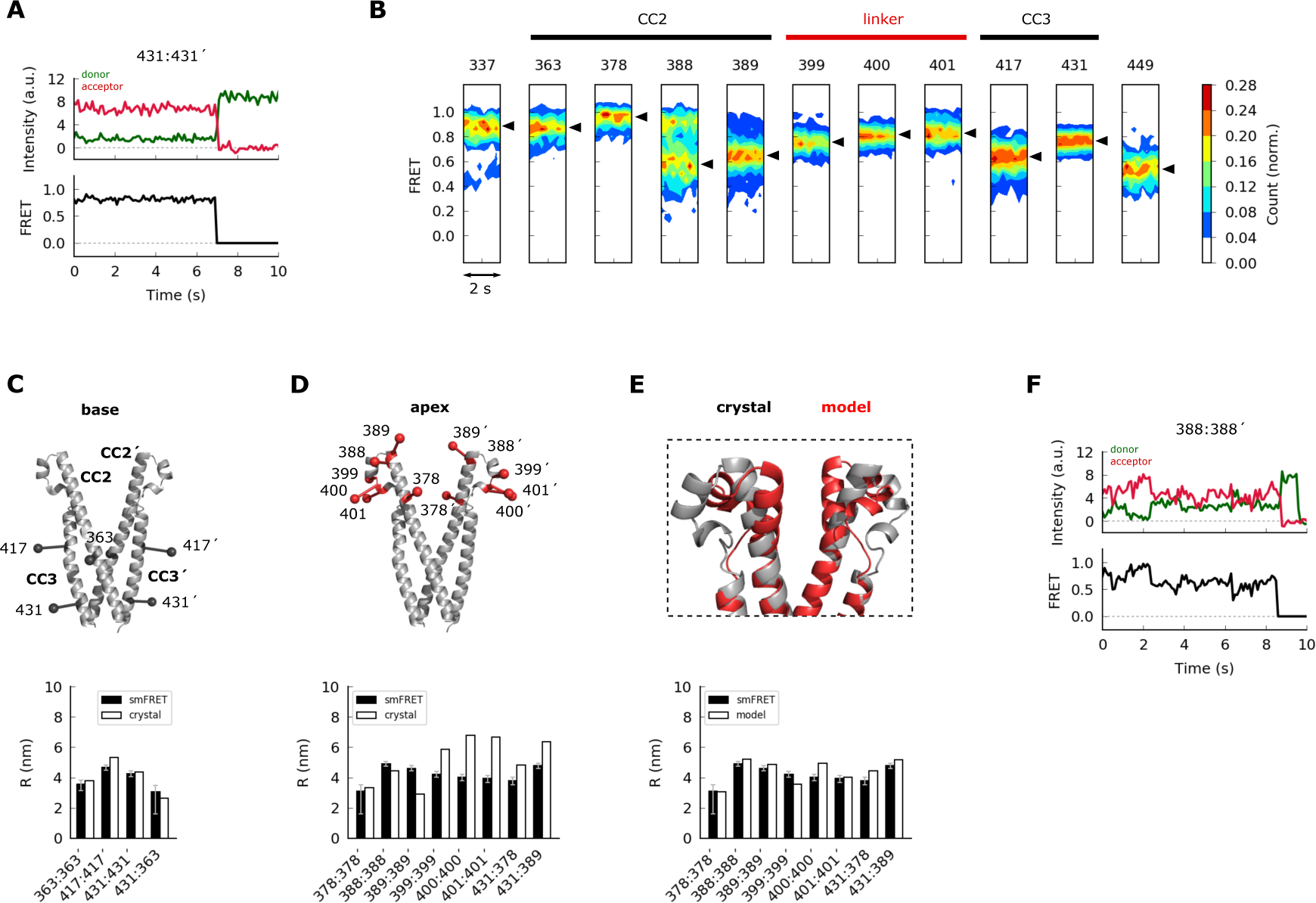
The apex of CAD in ctSTIM1 deviates from the CAD crystal structure. (**A**) Representative smFRET recording at the base of CAD (431:431’) showing stable, high FRET. (**B**) Ensemble density plots of the initial 2 s of inter-subunit smFRET recordings for sites throughout CAD. Predominant FRET levels are shown by arrowheads to the right of each plot. (**C-D**) Comparison of inter-fluorophore distances from smFRET and the CAD crystal structure. Simulated fluorophores were represented by the average position of a central atom, indicated here as ball-on-stick models (see Methods). For sites near the dimer interface (the ’base’ of CAD, **C**), the smFRET-derived distances (*black bars)* agree closely with the crystal structure (*white bars*). Measurements in the apex (**D**) deviated from the crystal structure. (**E**) Modified structure (*red*) generated by a smFRET-constrained optimization of the crystal structure (*gray*). Outward rotation of the distal CC2 region (aa 379-391) around G379 as a pivot point restored close correspondence between smFRET-derived distances and the crystal structure (*bottom*). (**F**) Representative example of smFRET fluctuations in the CAD apex (388:388’).

In contrast to the base of CAD, smFRET measurements of the distal CC2 and CC2-CC3 linker regions (the CAD ‘apex’) deviated from those indicated in the crystal structure (Figure 2D). This is particularly interesting as mutagenesis studies have identified this region as critical for Orai1 binding and activation (Calloway et al., 2010; Korzeniowski et al., 2010; Thompson et al., 2018; Butorac et al., 2019; Wang et al., 2014). To provide a rough indication of the extent of the deviation implied by smFRET, we generated an alternative model of the CAD apex based on the crystal structure by allowing rotation around G379 to bring the simulated FRET efficiencies in line with our experimental results. In the resulting model the distal CC2 regions were rotated outward, while the regions around aa 400 were pulled closer together, effectively compacting the apical ‘V-shape’ (Figure 2E and see Methods). Whereas inter-subunit smFRET at aa 399-401 appeared as a well-defined peak, FRET levels at the distal end of CC2 were broadly distributed and appeared multimodal, particularly at aa 388 (Figure 2B). Accordingly, smFRET measurements in this region displayed occasional fluctuations (Figure 2F), suggesting conformational flexibility that was not expected from the uninterrupted rigid helix of the crystal structure. The smFRET-derived model in Figure 2E thus likely represents just one of a number of potential apical conformations sampled by spontaneous fluctuations of the distal CC2 domain.

As a complementary structural test, we analyzed inter-subunit cysteine-cysteine (cys-cys) crosslinking upon oxidation by copper phenanthroline (CuP, see Methods). Cysteines introduced throughout the CAD apex formed inter-subunit crosslinks, detected as ctSTIM1 dimers on non-reducing SDS-PAGE gels (Supplementary Figure 2A). This result shows that opposing residues in the apical region at least transiently come within the ∼6 Å range for cys-cys crosslinking (Qin et al., 2015), again inconsistent with the CAD crystal structure. Moreover, crosslinking efficiencies were similar for three adjacent apical cysteine pairs (399:399’, 400:400’, and 401:401’), suggesting significant rotational flexibility beyond the temporal resolution of our smFRET recordings. Taken together, these data show that flexibility and spontaneous fluctuations cause the CAD apical region to deviate significantly from the crystal structure, unlike the stable basal region.

### CC1α1 docks parallel to CC3 in a domain-swapped configuration

The CC1 domain controls the activation of STIM1 *in vivo* by releasing CAD for interaction with Orai1 (Zhou et al., 2013; Ma et al., 2015). Although it is clear that the membrane-proximal CC1α1 domain can interact with CAD to inhibit STIM1 (Ma et al., 2015), their relative orientations remain undefined. We detected high inter-subunit FRET efficiencies between labels at the N-termini of CC1α1 and CC3 (red histogram in Figure 3A, top), as well as between labels at their C-termini (blue histogram in Figure 3A, bottom), indicating that the CC1α1 domain (a continuous α-helix (Cui et al., 2013; Rathner et al., 2020)) is oriented parallel to CC3 in the predominant ctSTIM1 conformation.

**Figure 3.**
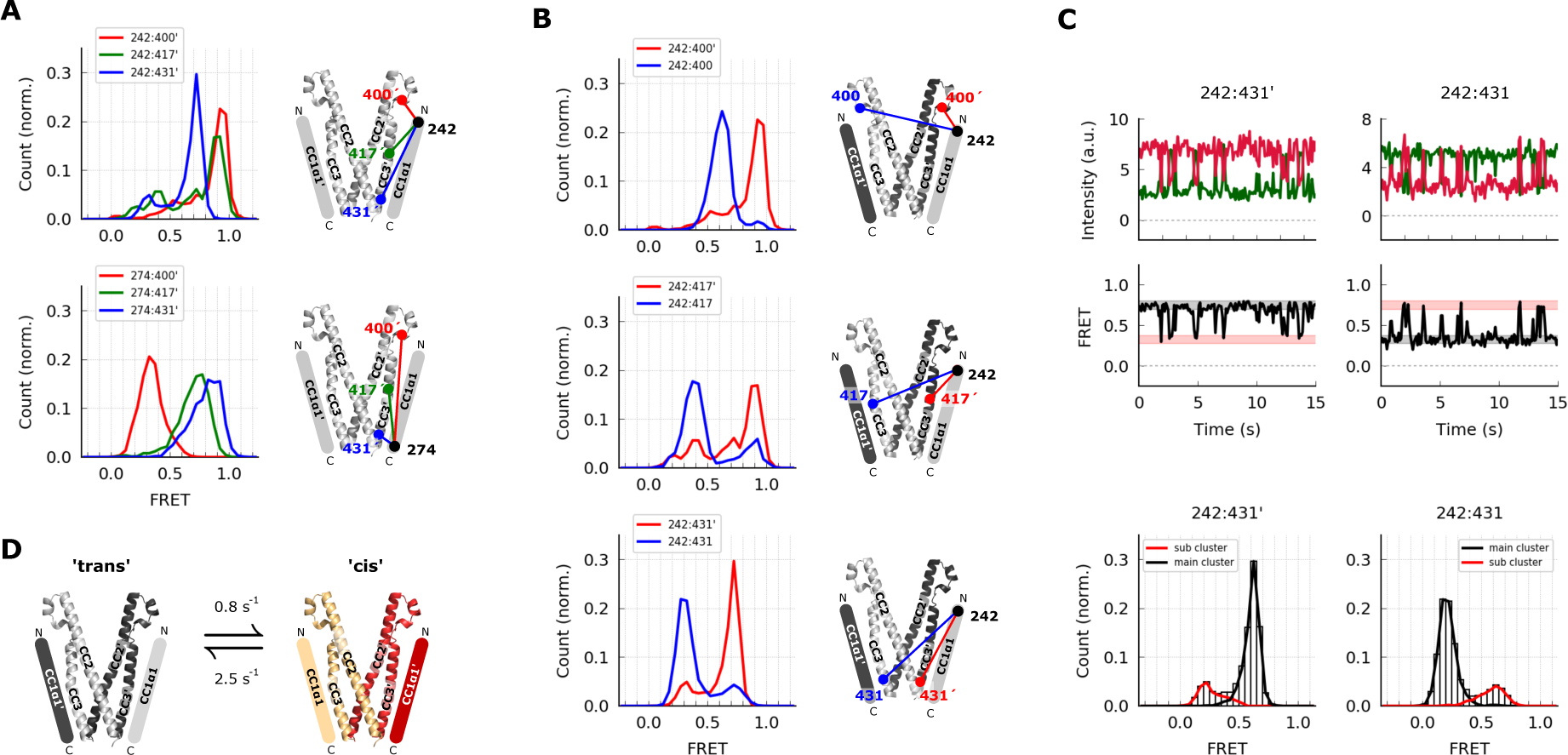
The CC1α1 domain is near and parallel to the CC3 domain of the opposing CAD subunit. (**A**) Asymmetric inter-subunit smFRET measurements between CC1α1 and CC3 reveal a parallel orientation, with high FRET levels at 242:400’ (*top, red*) and 274:431’ (*bottom, blue*) indicating close apposition. The crystal structure of CAD is shown with schematic representation of CC1α1 domains. (**B**) The predominant smFRET level between CC1α1 and CC3 was consistently higher for inter-subunit measurements (*red*) than intra-subunit measurements (*blue*). This pattern indicates a predominant domain-swapped configuration in which CC1α1 docks close to the CC3 domain of its opposing subunit. (**C**) Inter- and intra-subunit recordings (242:431’ and 242:431, respectively) displayed brief spontaneous smFRET transitions between two states indicated by red and black bars. In both cases fluctuations occurred between the same two FRET levels, suggesting the CC1α1 domains switched sides on CAD (see also Supplementary Figure 3A). (**D**) Schematic illustration of spontaneous conformational transitions of the CC1α1 domain. In the predominant conformation, CC1α1 is closely apposed and parallel to CC3 on its opposing subunit in a domain-swapped configuration (’trans’). Occasionally, CC1α1 switches to interact briefly with CC3 on its own subunit (’cis’). Rate constants were derived from dwell time analysis of smFRET traces (Supplementary Figure 3A).

To determine whether CC1α1 associates with the CAD of the same subunit or the adjacent one within the dimer, we compared inter-subunit and intra-subunit FRET efficiencies between labels at the N-terminus of CC1α1 and three sites in CAD (Figure 3B). The predominant inter-subunit FRET efficiencies were consistently higher than intra-subunit FRET efficiencies, demonstrating that in the predominant ctSTIM1 conformation, CC1α1 engages CAD in a domain-swapped configuration. We also observed spontaneous and reversible transitions from the predominant FRET state, in which FRET efficiencies fluctuated between the same two levels for both inter- and intra-subunit measurements (Figure 3C). Similar fluctuation behavior and transition kinetics were observed at all three CAD sites (Supplementary Figure 3A), suggesting a conformational transition in which the two CC1α1 domains briefly switch sides on CAD, probably using the same binding interface (Figure 3D).

### The CC1α3 domains are compactly folded and directed away from CAD

*In vivo*, mutations or deletions of the CC1α3 domain activate STIM1 (Korzeniowski et al., 2010; Yang et al., 2012) but the underlying mechanism, in particular the proposal that CC1α3 interacts directly with CAD (Yang et al., 2012), has been questioned (Zhou et al., 2013). We measured predominantly high inter-subunit FRET efficiencies of ∼0.9 between labels at the N- and C-terminal ends of the CC1α3 domains (aa 312 and 337, respectively), indicating that the two domains are closely apposed and parallel (Figure 4A). To determine their orientation relative to CAD, we measured intra-subunit FRET efficiencies between labels at either end of the CC1α3 domain to three sites in CC2. While FRET efficiency from the CC1α3 C-terminus to the proximal CC2 domain was high as expected (red histogram in Figure 4B, top), low FRET efficiencies were measured from the CC1α3 N-terminus, indicating that the N-terminus is directed away from CAD (Figure 4B, bottom).

**Figure 4.**
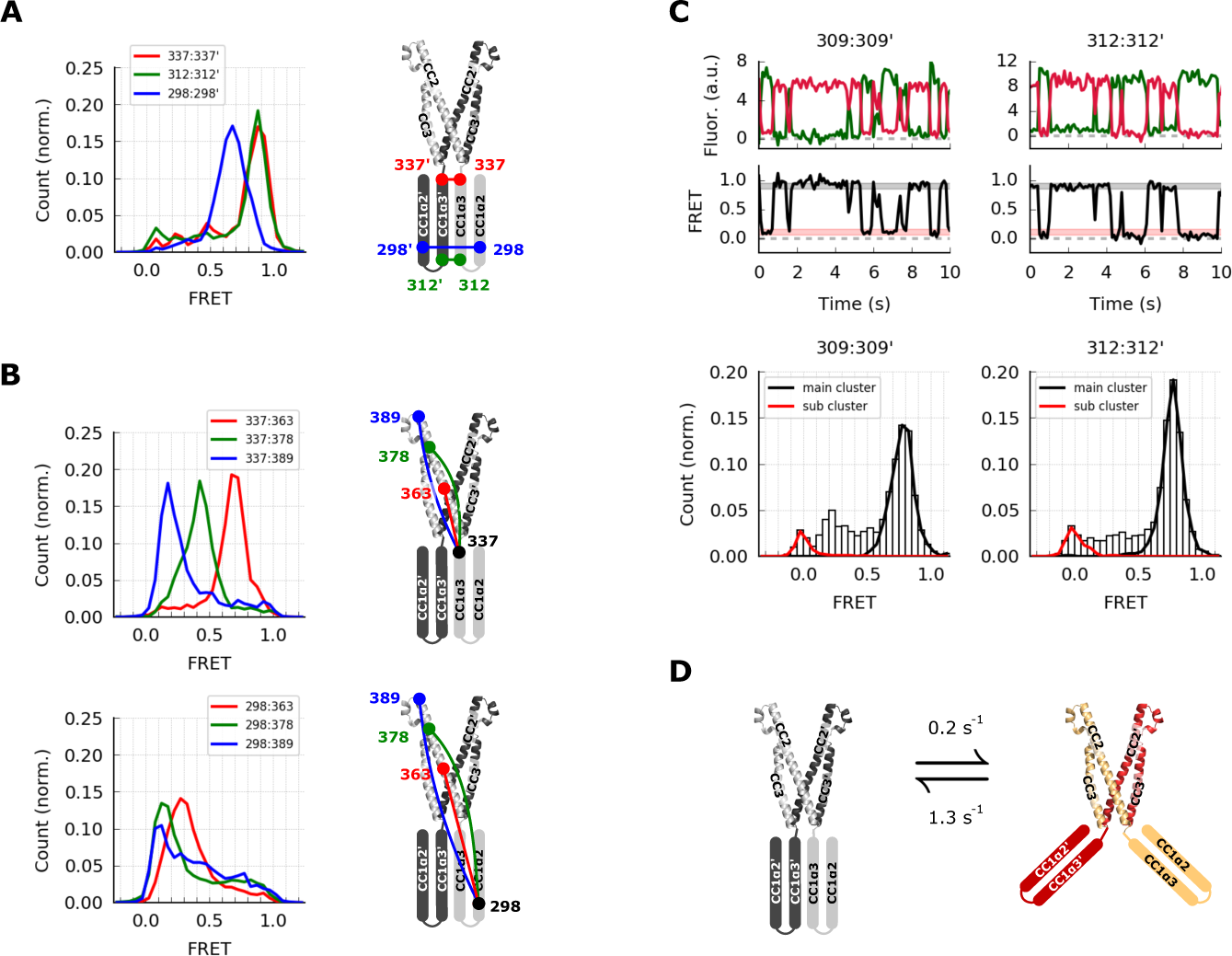
The CC1α2 and CC1α3 domains are closely apposed and directed away from CAD. (**A**) High inter-subunit smFRET at the N- and C-termini of the CC1α3 domain (aa 312 and 337, respectively) indicates close parallel apposition of the two helices. The crystal structure of CAD is shown with a schematic depiction of CC1α2 and CC1α3 domains. (**B**) (*top*) High intra-subunit smFRET from aa 337 to sites on CC2 declined progressively with distance, while intra-subunit smFRET from aa 298 (near the CC1α2/3 linker) to sites in CC2 was low (*bottom*), indicating that the CC1α2 and α3 domains are directed away from CAD. (**C**) Inter-subunit recordings near the CC1α3 N-terminus showed brief transitions from the predominant high- FRET state to a transient low-FRET state. Ensemble histograms show similar main and transient FRET states for neighboring sites aa 309 and 312. (**D**) Schematic depiction of spontaneous conformational transitions of the CC1α3 domains. CC1α3 domains are predominantly in close parallel apposition (*left*) and occasionally transition to a transient open configuration (*right*). Rate constants were derived from dwell time analysis of smFRET traces (Supplementary Figure 3B).

Stable high inter-subunit FRET efficiencies at the CC1α3 C-terminus (aa 337) were consistent with its proximity to the compact, stable CAD base. However, label sites near the N-terminus of CC1α3 displayed brief (0.8 s average lifetime), large transitions to a low FRET state, suggesting that the N-termini of CC1α3 intermittently splay out from a stable pivot point at the base of CAD (Figure 4C, D). While these large conformational transitions were most common, further analysis revealed additional substates in the smFRET trajectories (Supplementary Figure 3B, C).

### A model for the CC1-CAD complex

Through the integration of many inter-molecular distance measurements, smFRET can be applied to build models of tertiary protein structure (Brunger et al., 2011). We used 36 unique inter- and intra-subunit smFRET measurements to develop a structural model of the resting conformation of CC1 subdomains relative to the optimized model of CAD depicted in Figure 2E. Each CC1 domain in the ctSTIM1 dimer was modeled as a set of three rigid α-helical domains connected by flexible linkers, consistent with bioinformatic analysis (Soboloff et al., 2012), NMR measurements of the CC1 fragment (Rathner et al., 2020), and distances estimated from intra-subunit smFRET efficiencies (239:274, 274:307, and 307:337; Supplementary Figure 1).

We selected randomly generated models that met the criteria of satisfying smFRET-derived distance constraints while avoiding steric clashes with each other or with CAD (see Methods and Supplementary Figure 4A, C for details of the modeling process). Two classes of models emerged that were highly similar but distinguishable by the organization of the CC1α2/3 domains. In ∼70% of solutions the CC1α2/3 domains were stacked parallel (representative model in Figure 5A from an ensemble of models shown in Supplementary Figure 4Bi), while the remaining solutions had an alternative, wedged CC1α2/3 topology (Figure 5A and Supplementary Figure 4Bii). Importantly, both classes of solutions displayed the defining features described above, including closely apposed CC1α3 domains directed away from CAD and CC1α1 domains oriented parallel to CC3 in a domain-swapped configuration.

As an independent test of the smFRET-derived models, we applied the lysine crosslinking agent BS^3^ combined with mass spectrometry (XLMS) to identify residues in close proximity to each other (Rappsilber, 2011). BS^3^ XLMS confirmed key structural arrangements of both models of the CC1-CAD complex, including the compact packing of CC1α2/3 domains and the domain-swapped CC1α1-CAD interaction (Supplementary Figure 5A-E).

Although the models do not specify the orientation of side chains in the CC1 helices, they highlight two regions for potential inter-subunit coiled-coil interactions in the CC1-CAD complex: an antiparallel apposition of heptad repeats in CC1α2 and CC1α3’ (Figure 5B) and a parallel apposition of heptad repeats in CC1α1 and CC3’ (Figure 5C). The CC1α2/α3’ interface is reminiscent of the tight, antiparallel interaction of CC1α2 and CC1α3’ domains in the CC1 dimer crystal structure (Cui et al., 2013), an arrangement that was further supported by the formation of amine-carboxy crosslinks in ctSTIM1 by 1-ethyl-3-(3- dimethylaminopropyl)carbodiimide hydrochloride (EDC), a zero-length crosslinker (Figure 5D and Supplementary Figure 6).

Interestingly, the CC1α1:CC3’ interface in the smFRET-derived models juxtaposes multiple hydrophobic residues that have been previously implicated through mutagenesis in stabilizing the CC1-CAD complex: L248, L251, L258, L261 in CC1α1 and L416, V419, and L423 in CC3 (Fahrner et al., 2014; Muik et al., 2011; Zhou et al., 2013; Ma et al., 2015). To refine the CC1α1:CC3’ interface, we aligned the crystal structure of the CC1α1 domain (aa 246-271) to the smFRET-derived model and performed a computational docking simulation using Rosetta software. The resulting model indicates an extensive pairing of hydrophobic residues predicted to stabilize the CC1α1:CC3’ interface (Figure 5E), providing a plausible explanation for the activating effects of disruptive mutations at these sites.

### The Stormorken mutation R304W activates ctSTIM1 by releasing CC1α1 from CAD

The close packing of CC1α1 and CAD in our model suggests that ctSTIM1 is primarily inactive. Accordingly, when expressed with Orai1 in HEK293 cells, mCh-ctSTIM1 only slightly increased [Ca^2+^]i compared to mCh-CAD (Supplementary Figure 7), in agreement with previous evidence that ctSTIM1 is a relatively weak activator of Orai1 (Park et al., 2009; Korzeniowski et al., 2010; Yuan et al., 2009; Muik et al., 2009).

Deletion of the CC1 domain (mCh-ctSTIM1-ΔCC1) restored ctSTIM1 activity to the level seen with CAD, as did the introduction of R304W, a naturally occurring mutation that causes Stormorken syndrome by constitutively activating STIM1 (Misceo et al., 2014; Morin et al., 2014; Nesin et al., 2014) (Supplementary Figure 7). Interestingly, introduction of R304W into ctSTIM1 almost completely eliminated the high inter-subunit FRET between CC1α1 sites and the CC3 N-terminus *in vitro*, indicating the release of CC1α1 from CAD (Figure 6A, left and center). A more moderate decrease in FRET efficiency was observed relative to the CC3 C-terminus (aa 431), suggesting that CC1α1 pivoted around the CC1α1/2 linker (Figure 6A, right).

**Figure 6.**
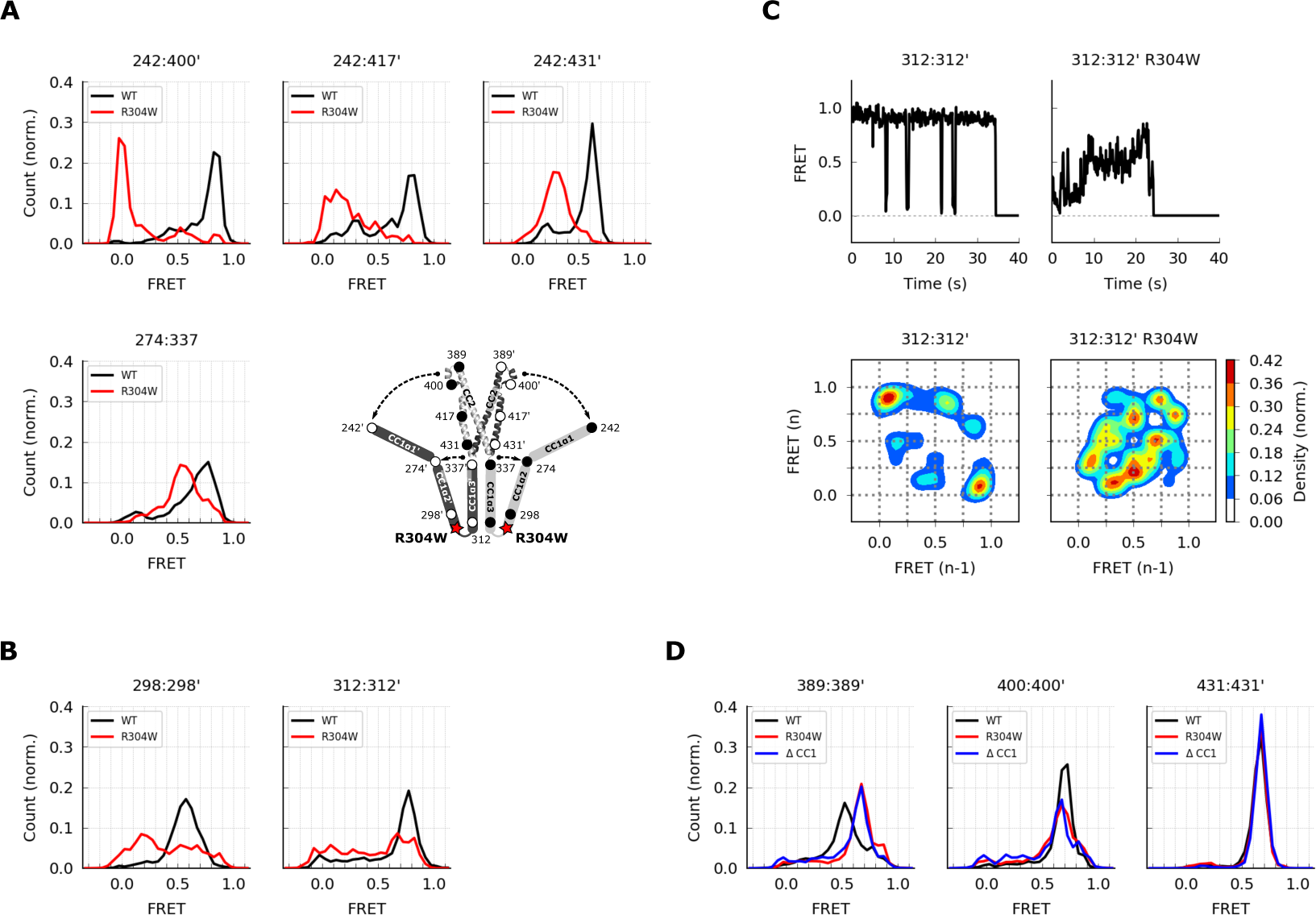
The Stormorken R304W mutation releases CC1 from CAD. (**A**) The R304W mutation reduced inter-subunit smFRET between aa 242 on CC1α1 and sites 400’, 417’, and 431’ on CC3’, indicating separation of CC1α1 from CAD (*top*). The mutation also reduced intra-subunit FRET for 274:337, suggesting a widening of the angle between CC1α2 and CC1α3 (*bottom*). (**B**) The predominant FRET levels for 298:298’ (*left*) and 312:312’ (*right*) were diminished by R304W, suggesting the compact parallel structure of CC1α2/3 domains was disrupted in favor of a dilated, low-FRET state. (**C**) Effects of R304W on inter-subunit FRET fluctuations at 312:312’, near the CC1α3 N-terminus. The dominant large-amplitude fluctuations of the wild-type (*top left*) were entirely absent in the mutant, which instead displayed frequent small fluctuations at a range of FRET levels (*top right*). The different fluctuation modes are summarized in ensemble transition density plots (see Methods) which show a single dominant FRET transition for wild-type molecules (*bottom left*, 38 molecules), and a multimodal density distribution for R304W mutants (*bottom right*, 25 molecules). (**D**) In the CAD apex, R304W or deletion of the entire CC1 region (ctSTIM1-ΔCC1) increased the FRET at 389:389’ (*left*) and reduced it at 400:400’ (*center*), suggesting that the apex interacts with CC1α1 in the WT conformation. Neither R304W nor ΔCC1 affected FRET at 431:431’ (*right*) in the CAD base.

To determine how R304W releases the CC1α1 domain given its distant location at the C-terminal end of CC1α2, we examined its effect on folding of the CC1α2 and CC1α3 domains.

Intra-subunit smFRET measurements indicated that R304W increases the separation between the CC1α2 N-terminus and the CC1α3 C-terminus (274:337) (Figure 6A, bottom left), suggesting it has opened the tight angle between CC1α2 and CC1α3. R304W also shifted CC1α2:CC1α2’ and CC1α3:CC1α3’ FRET to lower levels (Figure 6B), again consistent with opening of the compact inactive structure.

The R304W mutation also significantly altered the structural dynamics of CC1α2/α3. In the wild-type protein, the predominant inter-subunit FRET efficiency at the CC1α3 N-terminus was high, interrupted by occasional transitions to a low FRET state (Figure 4C and Figure 6C, left). In contrast, the R304W mutation appeared to destabilize the CC1α3 pair, such that it transitioned between high and low FRET through a series of intermediates states (Figure 6C, right). Taken together, our data reveal several effects of R304W on the CC1 domain: it destabilizes the tight packing of CC1α3 domains, causes CC1α2 to splay out from CC1α3, and completely releases CC1α1 from CAD.

We also assessed effects of R304W on the structure of CAD. While the R304W mutation or the deletion of CC1 (ΔCC1) did not affect inter-subunit FRET efficiencies at the base of CAD (431:431’), they did alter the smFRET distributions at the CAD apex (389:389’ and 400:400’) (Figure 6D). Together, these results suggest that the association of CC1α1 with CAD influences the conformation of the flexible apex without affecting the arrangement of helices at the more stable CAD base.

### The CC1 domains refold into a parallel dimer during STIM1 activation

We applied cysteine crosslinking to obtain complementary structural information for the CC1 domains. Inter-subunit cysteine crosslinking in ctSTIM1 was notably robust in the CC1α2/α3 linker (aa 307) and the C-terminus of CC1α3 (aa 339) (Supplementary Figure 8A). Disulfide formation at T307C prevented BS^3^ from crosslinking CC1α1 and CAD, indicating that T307C crosslinking is incompatible with the CC1α1-CAD inhibitory clamp (Supplementary Figure 5F). Similar findings were obtained for the activated ctSTIM1 mutants R304W and L248S/L251S (Supplementary Figure 5F), suggesting that close apposition and crosslinking at T307C resulted from spontaneous, transient visits to an activated state of ctSTIM1.

Under physiological conditions, full-length STIM1 (flSTIM1) in cells is activated by Ca^2+^ store depletion. To extend our *in vitro* ctSTIM1 results to flSTIM1, we examined cysteine crosslinking of inactive and active flSTIM1 *in vivo*. Cysteine mutations in flSTIM1 were made at selected sites throughout CC1, transiently expressed in HEK293 cells, and treated with the cell-permeant oxidizer diamide before or after store depletion induced by cyclopiazonic acid (CPA). Lysates of diamide-treated cells were then analyzed for the presence of crosslinked flSTIM1 dimers. Prior to store depletion, when flSTIM1 is inactive, little crosslinking was observed throughout CC1, with the exception of S339C near the junction with CAD (Figure 7A and 7b). In contrast, after store depletion prominent crosslinking occurred at A268C and T307C in addition to S339C (Figure 7A, B, Supplementary Figure 8). These results further support our conclusion above that T307C crosslinking of ctSTIM1 *in vitro* is enabled by transient activation events. More importantly, they suggest that activation *in vivo* brings all three CC1 subdomains together.

**Figure 7.**
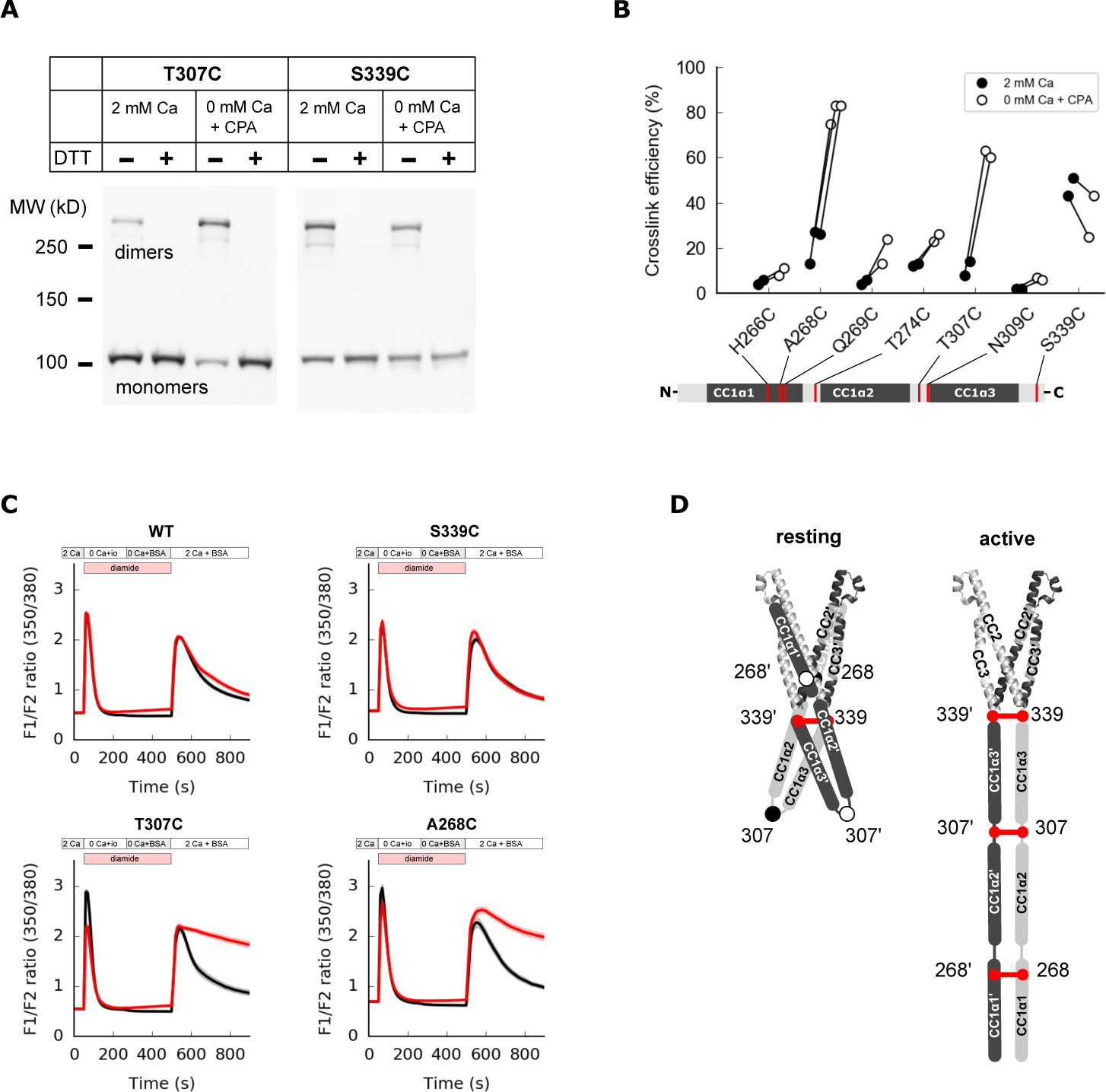
Close contacts form between CC1 domains along their entire length in the activated state of STIM1. (A) Full-length STIM1 (flSTIM1) mutants were transiently over- expressed in HEK293 cells for diamide-induced cysteine crosslinking *in vivo*. Western-blot analysis of cysteine crosslinking of flSTIM1-T307C (*left*) and flSTIM1-S339C (*right*), under resting (2 mM Ca^2+^) or store-depleted (0 mM Ca^2+^ + CPA) conditions. (**B**) Summary of flSTIM1 cysteine crosslinking before (*black*) and after (*white*) store depletion measured in individual paired experiments. While crosslinking at aa 268 and aa 307 strongly increased in the activated state, crosslinking at aa 339 occurred independent of STIM1 activation (see also **A** and Supplementary Figure 8B). (**C**) WT flSTIM1 and cysteine mutants were co-expressed with Orai1 for cytosolic calcium imaging (see Methods). For WT flSTIM1 cells, after store depletion by transient exposure to ionomycin (io), readdition of Ca^2+^ elevated [Ca^2+^]i due to SOCE, followed by a decline as SOCE deactivated from store refilling (*top left, black*). Diamide (*red*) did not affect the WT response. In contrast, diamide-induced crosslinking of T307C stabilized the active state, as evidenced by persistent calcium influx after ionomycin wash-out and store refilling (*bottom left, red*). Similar results were obtained for crosslinking A268C (*bottom right*, see also Supplementary Figure 8). Crosslinking of S339C did not inhibit deactivation of SOCE upon store refilling (*top right*). Each trace shows the mean and s.e.m. of the following numbers of cells (control/diamide) from at least two independent experiments: WT (91/106), S339C (44/41), T307C (38/46), A268C (31/47). (**D**) Schematic illustration of CC1 cysteine crosslinking in the resting (*left*) and activated (*right*) states of flSTIM1. In the resting conformation, CC1α1 and CC1α3 domains are kept apart, preventing crosslinking at locations upstream of aa 339. Upon store depletion, release of the CC1α1 domains from CAD promotes alignment of CC1 domains along their entire length, enabling crosslinking at aa 268 and 307.

To test this idea, we asked whether cysteine crosslinking after ER Ca^2+^ store depletion could lock flSTIM1 in the activated state, thereby preventing deactivation of STIM1 and SOCE upon store refilling. Ionomycin was added in Ca^2+^-free Ringer’s to deplete stores, after which ionomycin was removed and Ca^2+^ readded to allow Ca^2+^ influx through SOCE. In HEK cells expressing WT flSTIM1, [Ca^2+^]i increased rapidly upon Ca^2+^ readdition and subsequently declined due to store refilling and deactivation of STIM1 and Orai1, regardless of diamide treatment (Figure 7C). We obtained a similar result with flSTIM1-S339C, indicating that diamide-induced crosslinking at S339C does not impede flSTIM1 deactivation. In contrast, crosslinking at A268C and T307C prevented the normal deactivation of SOCE as indicated by the sustained elevation of [Ca^2+^]i after Ca^2+^ readdition. These findings suggest that all three CC1 subdomains are brought close together during STIM1 activation *in vivo* and remain closely associated in the final, Orai1-bound conformation (Figure 7D).

## Discussion

Precise regulation of SOCE is essential for many physiological processes and is achieved through control of transitions between inactive and active states of STIM1. These transitions are triggered by changes in ER [Ca^2+^] that toggle reversible interactions of the STIM1 CC1 domain with CAD. Through a combination of smFRET measurements and crosslinking assays, we have determined the arrangements of CC1 and CAD helices in inactive and active states of STIM1.

The results reveal how CC1 is oriented around CAD to stabilize the inactive state, and how its conformation changes upon activation to project CAD toward the PM to engage Orai1 and trigger SOCE.

In this study, we measured FRET efficiencies from single ctSTIM1 dimers to generate an overlapping set of intramolecular distance constraints for modeling tertiary structure (Choi et al., 2010; Brunger et al., 2011). Caution must be applied in using FRET signals to infer absolute distances, as FRET efficiency can also be affected by photophysical transitions or interactions with the local environment (Weiss, 1999, 2000). We employed several strategies to avoid misinterpretation of the data. First, by including a large number of different FRET pairs (36 pairs for CC1-CAD), no major feature of the CC1-CAD model was dependent on any single measurement, safeguarding against potential problems with individual label sites (Supplementary Figure 4A, C). Second, we independently verified the CC1-CAD model by mass spectrometry analysis of BS^3^-crosslinked protein (Schmidt and Robinson, 2014) (Supplementary Figure 5A- E). Third, measurements of key smFRET amplitudes and spontaneous structural transitions were confirmed at multiple nearby sites (Figure 2 and Supplementary Figure 3). Finally, we applied relatively conservative distance error limits (±1 nm) in constraining the CC1-CAD model. These strategies taken together help validate the tertiary structures represented in our models.

### The conformation of CAD in the context of ctSTIM1

smFRET measurements demonstrated that CAD within the context of ctSTIM1 resembles the CAD crystal structure rather than the NMR structure of CC1α3-CC2 (Figure 1). The base is stable and compact (Figure 2), as expected given the hydrophobic interactions and hydrogen bonding between the two CAD subunits in this region (Yang et al., 2012). However, the CAD apex appeared flexible and diverged from the crystal structure (Figure 2). These differences most likely arose from stabilization of an alternate apical conformation in the crystal by hydrogen bonds between the apex and adjacent CAD subunits (Supplementary Figure 1B) (Yang et al., 2012). The flexible structure of the CAD apex we observed is intriguing because mutagenesis and cysteine-crosslinking studies suggest this region interacts with Orai1 to initiate channel activation (Calloway et al., 2010; Korzeniowski et al., 2010; Thompson et al., 2018; Butorac et al., 2019; Wang et al., 2014), and conformational flexibility can facilitate protein-protein interactions through effects on the thermodynamics of binding (Grünberg et al., 2006).

Thus, it is likely that the CAD apex in complex with Orai adopts a structure different from that depicted in the CAD crystal and distinct from the NMR structure of CC1α3-CC2 (see below). Release of CC1α1 from CAD, or deletion of the CC1 domain altogether, did not cause a gross change in CAD structure, although it did alter the conformation of the CAD apex (Figure 6).

This raises the possibility that upon CC1α1 dissociation the apex changes shape, perhaps facilitating its binding to Orai1.

### Defining structural roles for all three CC1 subdomains in the STIM1 inhibitory clamp

Our results reveal how the three helical regions within CC1 collaborate to regulate the activity of STIM1. Prior evidence indicated that CC1 autoinhibits STIM1 in resting cells but left unanswered fundamental questions surrounding its structure. Korzeniowski and colleagues originally reported that neutralization of acidic residues in the CC1α3 domain activated STIM1 and proposed an ‘inhibitory clamp’ mechanism in which this region binds to a basic region within CAD to shield it from Orai in resting cells (Korzeniowski et al., 2010). This hypothesis potentially explained the activation of STIM1 by deletion of the CC1α3 domain (Yang et al., 2012). However, subsequent studies showed that CC1α3 alone is not sufficient to maintain the inactive state, as CC1α3-CAD is fully active (Zhou et al., 2013); moreover, intermolecular FRET between ER-tethered STIM1 fragments showed that CC1α1 can bind to CC3 (Fahrner et al., 2014), and ER-tethered CC1α1 by itself can bind CAD and keep it from binding to Orai1 in the PM (Fahrner et al., 2014; Ma et al., 2015). These findings and the activating effects of mutations in CC1α1 (Fahrner et al., 2014; Muik et al., 2011; Zhou et al., 2013; Ma et al., 2015) and CC3 (Muik et al., 2011) shifted the focus to a CC1α1-CC3 interaction as the basis for the inhibitory clamp, and the requirement for CC1α3 has remained unexplained. In addition to defining the CC1α1-CC3 interaction (below), our smFRET-derived structure can now clarify why deletion of CC1α3 disables the inhibitory clamp. CC1α2 and CC1α3 are compactly folded and directed away from CAD, allowing CC1α1 to engage CC3 (Figure 5); in the absence of CC1α3, the rigid CC1α2 helix would prevent CC1α1 from binding to CAD, which would then be free to interact with Orai1. Consistent with this explanation, deletion of CC1α2 in addition to CC1α3 allows the inhibitory clamp to function normally (Fahrner et al., 2014).

Based largely on mutagenesis studies, Zhou and colleagues proposed that the inhibitory clamp results from antiparallel binding of CC1α1 to CC3, involving interactions between L261 and L416 and between L258 and V419 (Ma et al., 2015). In contrast, our smFRET measurements show that CC1α1 binds parallel to CC3 (Figure 3). Moreover, our model depicts interactions of L248 and L251 (CC1α1) with L416 (CC3), and of L258 and L261 (CC1α1) with L423 and L427 (CC3) (Figure 5), which may explain the activating effects of non-conservative mutations at these sites (Fahrner et al., 2014; Muik et al., 2011; Zhou et al., 2013; Ma et al., 2015). Interestingly, our data reveal that CC1α1 preferentially associates with CC3 of the adjacent subunit in the CAD dimer (Figure 3). Such a domain-swapped configuration may enhance the stability of the inactive state through inter-subunit interactions and could promote cooperativity between subunits during structural transitions.

Consistent with a high degree of flexibility, CC1 displayed large spontaneous FRET fluctuations. The CC1α2/α3 domains occasionally transitioned from a compact, high-FRET resting state to a low-FRET open state (Figure 4), while the CC1α1 domains briefly switched sides on CAD, probably trading binding interfaces (Figure 3). Differences in dwell times suggest these two types of fluctuations occurred independently (Supplementary Figure 3), but further studies will be needed to determine how they relate to structural changes during activation of membrane-inserted flSTIM1. During the preparation of this manuscript, a report appeared describing a solution NMR structure of the isolated CC1 peptide, in which the CC1α1/α2/α3 helices form a monomeric three-helix bundle (Rathner et al., 2020). Rathner et al have proposed that this structure represents a pre-activated state of CC1 in which the interaction of CC1α1 and CC1α2 has destabilized the clamp formed by CC1α1 and CC3. Our smFRET experiments show that when CC1α1 associates with CC3 in ctSTIM1, the CC1α2 and CC1α3 domains primarily interact with each other to form a compact bundle pointing away from CAD (Figure 5); in the context of this recent report, we hypothesize that the fluctuations of CC1α2/α3 we observed may represent spontaneous transitions into such a pre-activated state.

The R304W Stormorken mutation activates ctSTIM1 (Supplementary Figure 7), consistent with its ability to activate flSTIM1 (Fahrner et al., 2018; Nesin et al., 2014; Morin et al., 2014; Misceo et al., 2014). smFRET measurements show that R304W destabilizes the compact packing of CC1α2/3 domains, increasing the angle between CC1α2 and CC1α3 helices and fully releasing CC1α1 from CAD (Figure 6). These results are consistent with NMR studies showing that R304W causes N-terminal extension of the CC1α3 helix, thereby stiffening the CC1α2/α3 linker (Fahrner et al., 2018; Rathner et al., 2020).

### A structural view of quiescent STIM1 and its transition to the activated state

Our results provide a first view of how the CC1 domain of STIM1 sequesters CAD near the ER membrane in resting cells to prevent it from interacting with Orai. The parallel arrangement of CC1α1 and CC3 in our smFRET-derived structure predicts that the CAD apex of inactive STIM1 is juxtaposed to the ER membrane and pointed away from the PM *in vivo* (Figure 8A). This orientation implies that to activate STIM1 and SOCE, the CAD apex must rotate through a large angle (up to 180°) and translocate more than 10 nm towards the PM - a massive conformational change reminiscent of the refolding and extension of viral envelope proteins like hemagglutinin when stimulated to trigger membrane fusion (Harrison, 2008). Hirve et al. showed that activation of the flSTIM1 cytosolic domain is initiated by formation of a coiled-coil in the TM domains and the proximal part of CC1α1 up to L251 (Hirve et al., 2018). As an extension to these findings, we found that after store depletion, inter-subunit disulfide bonds form at A268C (C-terminus of CC1α1) and T307C and S339C (N- and C-termini of CC1α3, respectively; Figure 7 and Supplementary Figure 8). Taken together, these results suggest that all three helical domains in CC1 are closely paired in the active dimer, which would effectively move CAD across the ∼15 nm gap of the ER-PM junction to reach Orai (Wu et al., 2006).

**Figure 8.**
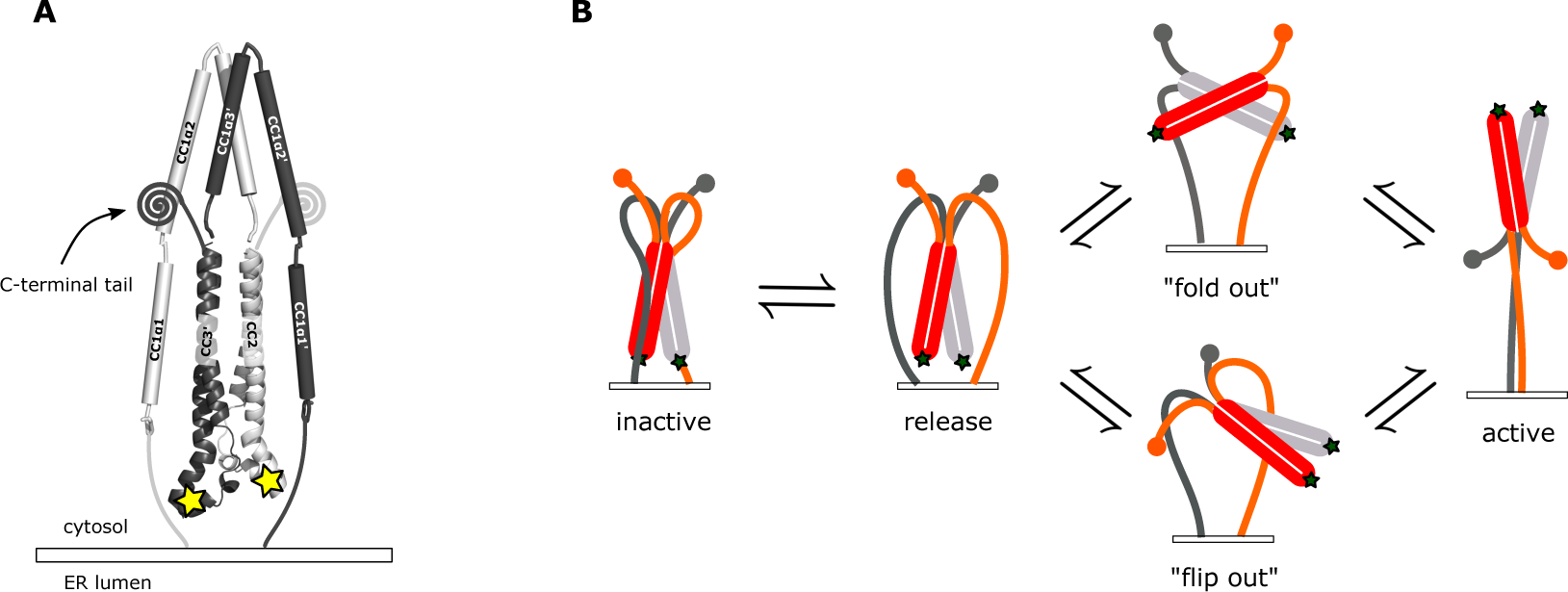
Possible conformational trajectories of flSTIM1 activation *in vivo*. (**A**) In the resting state, the parallel orientation of CC1α1 and CC3 domains implies that CAD is held with its apex close to and pointed towards the ER membrane, effectively shielding critical interaction sites (*stars*) from engaging with Orai. The region C-terminal to CAD is abbreviated as its conformation is not known. (**B**) Alternative models for transitions to the active state of flSTIM1. In the “fold out” model (*top*), following the release of CC1α1, CAD undergoes a symmetric conformational change in which the CC2 domains pass through a transient antiparallel state, providing a free path for the region downstream of CAD to extend towards the plasma membrane. In the “flip out” model (*bottom*), CAD maintains its parallel resting conformation and undergoes an asymmetric outward rotation from between the CC1 domains. Assuming that the initial resting conformation is symmetric, one of its C-terminal domains (*gray*) would have to be carried through the gap between the two CC1 domains.

Our CC1-CAD model specifies the possible types of conformational changes that must occur to activate flSTIM1 and present the CAD apex to Orai (Figure 8B). We can envision two potential solutions. In a ‘flip-out’ mechanism, CAD rotates as a whole by 180°. Although our smFRET data indeed indicate that release of CC1α1 does not cause a gross conformational change in CAD (Figure 6), this kind of CAD rotation would imply a complex asymmetric intermediate, in which one of the C-terminal domains (aa 449-685) downstream of CAD would need to be threaded between the two CC1 domains. Alternatively, in a ‘fold-out’ model, CAD undergoes a symmetric conformational change while moving its apex away from the ER membrane. This model posits that during activation the CC2 domains rotate to transiently assume an antiparallel state, reminiscent of the NMR structure (Figure 1A and 8B). It is important to note that our cysteine crosslinking results indicate the CC1α3 C-termini are closely apposed in the Orai-bound state (Figure 7D, S339C), which is incompatible with the splayed configuration of CC1α3 domains in the NMR structure. Thus, if the antiparallel conformation exists it is likely to represent a transient intermediate during the activation of STIM1 (Stathopulos et al., 2013).

The ultimate endpoint in delineating the mechanism of SOCE is an atomic-level description of the pathway of STIM1 activation by store depletion. To reach this goal the starting, intermediate, and final conformations of STIM1 after store depletion must be resolved, as well as the structure of the activated STIM-Orai complex. While approaches like cryo-EM certainly offer an effective strategy for determining structures, time-resolved measurements such as those afforded by smFRET will be essential to identify structural transitions in the activation pathway. The results obtained thus far through smFRET of ctSTIM1 have revealed critical measurement sites that offer a starting point for these studies. Ultimately, the identification of transient intermediate STIM1 conformations and rate-limiting steps in the activation process may reveal critical transitions that could be targeted to manipulate SOCE for therapeutic benefit.

## Materials and methods

### DNA constructs

For samples that were encapsulated in liposomes for surface immobilization, DNA encoding ctSTIM1 (aa 233-685) was amplified by PCR from full-length human STIM1 (Origene), appending an N-terminal NcoI cleavage site, and a C-terminal TEV protease recognition sequence (SENLYFQG) followed by a HindIII cleavage site. The ctSTIM1 insert contained a silent T1764C mutation in H588 to remove an endogenous NcoI site, and a G1310C mutation to replace the endogenous cysteine (C437S). ctSTIM1 inserts were ligated into the pET28a vector which encoded a C-terminal 6-His tag. ctSTIM1 cysteine mutants were created by site-directed mutagenesis (QuickChange II, Agilent). The translated ctSTIM1 protein retained the sequence SENLYFQ at the C terminus after cleavage of the 6-His tag by TEV.

For samples that were directly attached to a PEG-coated surface, the pAC6 vector was used with a C-terminal AviTag™ sequence GLNDIFEAQKIEWHE (Avidity). ctSTIM1 was first ligated into a pTEV5 vector which encoded a TEV-cleavable N-terminal 6-His tag. DNA encoding the 6-His tag, TEV recognition site and ctSTIM1 was then amplified by PCR, appending an XhoI cleavage site to the N terminus, and a HindIII cleavage site to the C terminus for ligation into the pAC6 vector. The resulting translated ctSTIM1 protein retained the N-terminal sequence GAS after TEV cleavage of the 6-His tag, and it had the AviTag™ sequence following a spacer KLPAGG on the C terminus. All constructs were verified by DNA sequencing.

### Protein expression, purification, and labeling

ctSTIM1 protein was expressed in *E. coli* based on standard methods described previously (Choi et al., 2010). Briefly, plasmid containing the ctSTIM1 insert was transformed to BL21 competent cells (New England Biolabs #C25271) for pET28a plasmids, or CVB101 competent cells (Avidity #CVB101) for pAC6 plasmids. Protein expression was induced with IPTG at OD ∼0.4 in 450 ml Luria Broth (LB), and for pAC6 plasmids biotin was added to a concentration of 50 µM. Bacteria were spun down and then resuspended in denaturing PBS (8 M urea, 100 mM NaH2PO4, 10 mM Tris (pH 7.4 with NaOH)) and lysed by sonication. TCEP (Thermo Fisher Scientific #77720) was added to a concentration of 0.5 mM. For intra-subunit smFRET samples, double-cysteine ctSTIM1 was mixed with cysteine-free ctSTIM1 in a 1:5 ratio to allow formation of ctSTIM1 heterodimers. Samples were purified by NiNTA purification (Qiagen #30210), and subsequently slowly dialysed using SnakeSkin™ (Life Technologies #68700) to 20/50 TBS (50 mM NaCl, 20 mM Tris (pH 7.4 with HCl)) containing 1 mM DTT. TEV protease (MCLAB #TEV-200) was added during dialysis at 10 μg/ml. Cleavage of the 6-His tag was usually almost 100%, as checked on SDS page gel. Samples were further purified by ion exchange on a HiTrap Q column (GE Healthcare Life Sciences #17-1153-01) and eluted in 250 μl aliquots of 20/250 TBS. All buffers contained 0.5 mM TCEP. Resulting ctSTIM1 samples were >90% pure and displayed the expected molecular weight (∼52 kD) by SDS-PAGE. Dimer concentration was typically 25-50 μM, measured by absorption photometry (NanoDrop 2000, ThermoFisher Scientific).

Protein labeling was performed as described previously (Choi et al., 2010) using Alexa Fluor 555 as donor fluorophore and Alexa Fluor 647 as acceptor fluorophore (Invitrogen Life Technologies #A-20346 and #A20347). For symmetric inter-subunit smFRET and for intra-subunit smFRET, donor and acceptor fluorophores were mixed in a 1:1 ratio before adding to the protein. For asymmetric inter-subunit smFRET, two different ctSTIM1 cysteine mutants were mixed separately with donor or acceptor fluorophores, to be recombined afterward into ctSTIM1 heterodimers. Labeling efficiency assessed by absorption photometry was typically ∼70% but varied for different locations; cysteine mutants with labeling efficiency below 50% were not used for smFRET experiments.

For recombination of asymmetric inter-subunit samples, labeled ctSTIM1 homodimers were denatured by adding urea to a concentration of 8 M. Donor and acceptor species were then mixed in a 1:1 ratio, and refolding was allowed to take place during slow dialysis as described above, resulting in a mixture containing 50% ctSTIM1 heterodimers. ctSTIM1 heterodimers with a single donor and a single acceptor fluorophore were individually selected during posthoc analysis of the smFRET experiments.

### Sample preparation

Flow cells for single-molecule fluorescence imaging were prepared according to established protocols (Roy et al., 2008). Briefly, strips of double-sided tape were put on a quartz microscope slide (Finkenbeiner) to form channel walls, with holes at the ends of each channel to create entry and exit points for buffer solutions. A microscope coverslip (Erie Scientific #24X40 1.5 001) was pressed on the tape strips and edges of the channels were sealed with two-component epoxy glue. For liposome experiments channels were coated with BSA-biotin (Sigma Aldrich #A8549). For direct surface attachment microscope slides and coverslips were coated with a 100:1 PEG/PEG-biotin mixture (Laysan Bio #BIO-PEG-SVA-5K & MPEG-SVA-5K) prior to flow cell construction. Neutravidin (Thermo Fisher CWA #31000) was flushed into the channels before loading the protein samples. 20/150 TBS (150 mM NaCl, 20 mM Tris, pH 7.4 with HCl) was used for rinsing in between steps.

For liposome encapsulation of fluorophore-labeled protein samples, biotinylated lipids (18:1 Biotinyl Cap PE, Avanti Polar Lipids #870273C) were mixed with egg-PC lipids (Avanti Polar Lipids #840051C) in a 1:100 ratio. Lipids were suspended in sample buffer and the mixture was passed back-and-forth (Extruder Set, Avanti Polar Lipids #610023) through a liposome extrusion membrane (PC Membranes 0.1μm, Avanti Polar Lipids #610005-1EA).

Liposomes were separated from free protein on a Sepharose® CL-4B column (Sigma Aldrich #CL4B200). The conditions resulted in on average <<1 ctSTIM1 dimer per liposome.

For direct surface attachment, biotinylated ctSTIM1 samples were diluted to a concentration of ∼100 pM and flushed into a PEG-coated, neutravidin-treated channel. Immediately prior to fluorescence imaging, channels were filled with 20/150 TBS containing 100 μM cyclooctatetraene (Sigma Aldrich #138924) (COT) and an oxygen scavenging system consisting of 1% D-glucose, 1 mg/ml glucose oxidase (Sigma Aldrich #G2133), and 0.04 mg/ml catalase (Sigma Aldrich #C9322).

### TIRF microscopy and smFRET measurements

Single-molecule FRET recordings were performed using a home-built through-the-objective total internal reflection fluorescence (TIRF) imaging system based on a Zeiss Axiovert S100 TV microscope. Alexa Fluor 555 (donor) and Alexa Fluor 647 (acceptor) fluorophores were excited by 532 and 637 nm wavelength lasers (OBIS 532 nm LS 150 mW, Coherent #1280719 and OBIS 637 nm LX 140 mW, Coherent #1196626). A Fluar 100x 1.45 NA oil-immersion objective (Zeiss) was used to establish TIR excitation and collect photon emission. Donor and acceptor emission were separated by a dichroic beamsplitter (Semrock FF652-Di01-25x36) and passed through band-pass filters (Semrock FF01-580/60-25-D for donor emission and Semrock FF01-731/137-25 for acceptor emission) mounted in an OptoSplit-II (Cairn Research), projecting the images side-by-side onto an EM-CCD camera (iXon DU897E, Andor). Hardware and image acquisition were controlled by BeanShell scripts from μManager (Edelstein et al., 2010), and image sequences were stored as stacked TIFF files.

Prior to image acquisition, the channel surface was scanned manually using low-intensity donor excitation to identify a suitable location for smFRET imaging, where the density of fluorescent spots appeared homogeneous and without contamination or aggregations. Donor and acceptor emission were then recorded in response to donor excitation, typically for 60 s, immediately followed by 1 s of acceptor excitation to directly identify acceptor fluorophores.

Images were acquired in frame-transfer mode with a 100-ms integration time, and laser intensity was set to obtain a trade-off between good signal-to-noise ratio and sufficient active time for both fluorophores (ideally tens of seconds), while inducing fluorophore bleaching within the recording time window. Typically, 10-20 movies were recorded in each of four channels per microscope slide. Measurements were performed at room temperature.

### smFRET data analysis

Fluorescence movies were analyzed using custom Python scripts. Donor and acceptor images were aligned using pre-recorded registration images, and fluorescing molecules were identified in the donor and acceptor channels by detecting local signal maxima within a five-pixel-diameter neighborhood. Donor signals that could not be matched to any acceptor signal were discarded.

Pixel values for each molecule were summed, and signals from sequential movie frames were stacked to construct the fluorescence time series for each molecule. The local background for each molecule was determined by calculating the median background value within a circular 35- pixel-diameter region in each frame and subtracting it from the raw fluorescence signal.

Leakage of donor signal into the acceptor channel in our system was approximately seven percent of the measured donor signal and was systematically subtracted from the measured acceptor signal.

γ correction was used to correct for differences in photon emission and detection probabilities of donor and acceptor fluorophores. The value of γ was empirically determined for each individual molecule (McCann et al., 2010), and the correction was applied by multiplying the donor fluorescence signal by γ before calculating the smFRET ratio. The average value from all 12,252 molecules was γ = 1.1.

Traces were selected for final analysis if all of the following criteria were met: (1) the signal-to-noise ratio was equal or greater than five; (2) the acceptor bleached in a single step before the donor bleached; (3) the γ factor was between 0.5 and 2.5; (4) any spontaneous fluctuations of donor and acceptor fluorescence were negatively correlated; and (5) if a donor bleach event was recorded, it occurred in a single step.

The FRET ratio E was calculated for each point in the donor and acceptor fluorescence traces (respectively ID and IA) where donor and acceptor were both active as E = IA/(IA + γID). A FRET histogram was constructed for each trace by distributing FRET amplitudes into 50 bins in a [-0.25, 1.25] range. Histograms were normalized by dividing each bin by the total number of FRET points in the trace. An ensemble histogram for multiple molecules was constructed by summing the normalized histograms of the individual traces and dividing each bin by the total number of molecules.

The part of the fluorescence traces where both donor and acceptor were active were parameterized by fitting a Hidden Markov model (HMM), using routines from the SMART software package (Greenfeld et al., 2012). To determine the optimal kinetic model, the Bayesian information criterium (BIC) was used to select between models with different numbers of states. To compare FRET levels between different molecules in an ensemble, each HMM state was assigned to a cluster based on its mean FRET value. Clustering was performed based on all observed states in the ensemble, using a k-means algorithm with a predetermined number of target clusters. The number of clusters was determined ad hoc depending on the shape of the FRET histogram. For each cluster the peak of its histogram was reported as its representative FRET level. The predominant FRET cluster was the cluster containing most FRET measurement points.

Periods between cluster transitions were used for constructing dwell time histograms.

The first and last period of each trace were discarded, so that traces with fewer than two transitions were not included in the dwell time analysis. For transition density plots each transition was described by a coordinate given by the FRET level just before the transition (horizontal axis) and the level after the transition (vertical axis). The local transition density was then calculated on a 50x50 grid ranging from FRET -0.25 to 1.25, by convolution with a Gaussian kernel with two-pixel standard deviation. To prevent individual molecules with many transitions from dominating the plot, each molecule contributed only one data point for each kind of transition. In this way the transition plot reports the number of molecules in which a certain kind of transition appeared.

For each protein sample, single-molecule recordings from multiple experiments were pooled to construct smFRET amplitude histograms. The total number of molecules sampled for each histogram (see Supplementary Figure 1) was determined such that additional molecules did not alter the position of the predominant peak in the histogram. Key smFRET results were replicated by multiple authors (biological replications with newly purified protein samples).

Molecules that did not produce smFRET were excluded, as they could not be properly interpreted.

### Molecular modeling and simulation

Molecular dynamics simulations and model optimization procedures were adapted from published procedures (Choi et al., 2010), and performed using Crystallography and NMR System (CNS) (Brunger et al., 1998; Brunger, 2007). Structures were visualized and manipulated using PyMOL (Version 1.8, Schrödinger, LLC.) and ProDy (Bakan et al., 2011). To predict distances between fluorophores, we performed molecular dynamics simulations using atomic models of donor and acceptor fluorophores. Because the structures of Alexa Fluor 555 and Alexa Fluor 647 are not available, we used the structures of the spectrally similar Cy3 and Cy5. Residues of interest on the STIM structure were mutated to cysteine and fluorophore models were covalently attached by CNS. While all protein atoms were held fixed in space, a simulated-annealing protocol sampled possible fluorophore conformations until a local energy minimum was reached. This procedure was repeated 100 times to obtain a distribution of fluorophore orientations, and the average location of the fluorophore center (taken as the location of the CAO atom) was computed from the resulting coordinates. When attached to sites in the CAD crystal structure the overall mean dye center protruded from the Cα atom of the attachment residue by 0.99 ± 0.08 nm for Cy3 and 1.10 ± 0.08 nm for Cy5. Since labeling was stochastic, the donor or acceptor could be on either side in a pair of label sites. To account for this, we averaged Cy3 and Cy5 center coordinates for each residue into a single “effective” center location. Fluorophores were represented in the simulations by a pseudo-atom positioned at this effective center location, and inter-fluorophore distances were then calculated as the distance between two pseudo-atoms.

According to Förster theory the distance between donor and acceptor fluorophores (*R*) can be derived from experimentally determined smFRET efficiency (*E*) as R = R0 (1/E - 1)^1/6^, where R0 is a proportionality factor reflecting the inter-fluorophore distance at which E = 0.5 (ref (Stryer and Haugland, 1967)). For Alexa Fluor 555 and Alexa Fluor 647 in water, the theoretical value of R0 is 5.1 nm, assuming fast isotropic re-orientation of the fluorophores. We used this value for calculating smFRET-derived distances.

For structural modeling of the CAD apex, we first created a symmetric version of the CAD crystal structure, as the original is slightly asymmetric in the apical region. We copied one subunit and aligned the proximal CC2 (aa 345-378) and CC3 (aa 408-436) domains to the adjacent subunit in the dimer, resulting in a symmetric pair. We then used CNS to allow the apex to relax to a new conformation, guided by smFRET-derived distance constraints (inter-subunit constraints at aa 388, 389, 399, 400, and 401, and intra-subunit constraint 431:389 on both subunits). Helices were treated as rigid bodies, while unstructured regions were treated as flexible chains with variable torsion angles. The distal CC2 helix (aa 379-391) was allowed to rotate around residue G379. During the relaxation of the apex, the proximal CC2 and CC3 domains were held fixed in space.

To reconstruct the arrangement of CC1 helices around CAD, we used the symmetric CAD structure with smFRET-optimized apical region and modeled CC1 as a chain of dimensionless nodes using custom Python code. Each node on the chain represented the center of a CC1 residue. CC1 chains were generated starting from aa 344 at the CAD N-terminus, by progressively connecting new nodes until the final residue aa 233 was reached. In unstructured regions, nodes were spaced every 0.38 nm and were generated at random angles ≥90°. Alpha- helical regions were modeled as a straight chain of nodes at 0.15 nm intervals. To create a symmetric pair of CC1 chains, nodes were generated for one subunit, and then copied and mirrored in CAD’s symmetry planes to create the chain for the other subunit. Each node was subjected to two criteria: (1) it cannot overlap with any residues in CC1 or CAD (volume exclusion in a 0.5 nm-radius sphere), and (2) it must obey smFRET-derived distance (DFRET) constraints. Fluorophores positions were not explicitly modeled on the CC1 chains but were accounted for by setting CC1:CC1’ distance constraints to the range of DFRET to DFRET - 2 nm, due to fluorophores extending up to 1 nm on both sides of the helix backbone. Because fluorophores were explicitly modeled in CAD, CC1:CAD distance constraints of DFRET to DFRET - 1 nm were applied. In cases of very high or very low FRET the margins were occasionally relaxed to account for the lack of sensitivity of the smFRET method at these extremes (see Supplementary Figure 4A for a table of all distance constraints used for the CC1:CAD model). If both criteria were met, the node was retained; otherwise it was rejected, and a new node was generated. If no suitable node was found, a new search was started for the preceding node. This process continued until the final N-terminal residue of CC1 (aa 233) was reached, and the resulting model was then stored. Solutions obtained in this way together defined a solution space within which CC1 obeyed the imposed constraints. We used the average of 50 solutions as a model for presentation (Figure 5 and Supplementary Figure 4).

To obtain a more detailed picture for the molecular interface of the CC1α1-CAD complex, we aligned the crystal structure of the CC1α1 helical segment (aa 246-271) to the smFRET-derived CC1 model, directing the hydrophobic sidechains to point toward CAD.

Residue A419 in the CAD crystal structure was mutated back to the original valine using Pymol. The CC1α1 helix was then docked onto CC3 using the flexpepdock server of Rosetta (London et al., 2011), which performed a molecular dynamics simulation with a fully flexible backbone. An initial docking was performed with distance constraints (2.5 ± 2.5 Å) between CG atoms of residue pairs L248:L416, L251:L416, L258:L423, and L261:L427, which were directly apposed in the smFRET-derived model. This was followed by an unconstrained simulation to obtain the final model (Figure 5E).

### Chemical crosslinking and mass spectrometry

For BS^3^ crosslinking of ctSTIM1, buffer was changed from Tris- to HEPES-based (150 mM NaCl, 20 mM HEPES (pH 8.0 with NaOH)) during the ion exchange procedure. ctSTIM1 samples at 10 μM (∼1 mg/ml) were allowed to incubate with 0, 20 and 50 μM BS^3^ (BS^3^-d0, Sigma Aldrich #21590 or BS^3^-d4, Sigma Aldrich #21595) for 30 min at room temperature. The reaction was then quenched for 15 min at room temperature with 20 mM Tris.

For crosslinking with EDC, the buffer was changed from Tris- to MES-based (250 mM NaCl, 100 mM MES, pH 6.0 with HCl) during the ion exchange procedure. To increase crosslinking efficiency, EDC (Sigma Aldrich #77149) was used in combination with sulfo-NHS (Sigma Aldrich #24520) to convert carboxyl groups into stable amine-reactive sulfo-NHS esters. ctSTIM1 samples at 100 μM (∼10 mg/ml) were allowed to incubate with 0/0, 20/62, 50/156 and 100/312 mM EDC/sulfo-NHS for 15 min at room temperature. Two volumes of PBS (250 mM NaCl, 100 mM Na2HPO4, pH 8.5 with NaOH) with 25 mM BME (pH 8.5) were then slowly added to quench the EDC and increase the pH for the subsequent amine reaction, incubated 2 h at room temperature and quenched with 40 mM Tris.

Crosslinked ctSTIM1 samples were run on SDS-PAGE gels, and monomer and dimer bands were cut and analyzed separately by mass spectrometry. For comparative BS^3^ crosslinking experiments, BS^3^-d0- and BS^3^-d4-treated samples were first mixed 1:1 before electrophoresis. Gel bands at 20 and 50 μM BS^3^ were combined for analysis (Supplementary Figure 5). For EDC experiments, the monomer band at 10 mM EDC and the combined dimer bands at 2, 5, and 10 mM were used (Supplementary Figure 6).

Mass spectrometry analysis of cross-linked samples was performed as described previously (Komolov et al., 2017). Briefly, gel bands were diced into 1x1 mm squares, reduced in 5 mM DTT at 55⁰C for 30 min, and then alkylated with 10 mM acrylamide for 30 min at room temperature to cap uncrosslinked cysteines. Following alkylation of free cysteines and washing of gel pieces, proteolytic digestion was completed using trypsin/LysC protease (Promega) in the presence of ProteaseMax (Promega) overnight at 37⁰C. Peptides were then extracted and dried under SpeedVac prior to LC/MS analysis.

Mass spectrometry experiments were performed using an Orbitrap Fusion Tribrid mass spectrometer (Thermo Scientific, San Jose, CA) with an Acquity M-Class UPLC system (Waters Corporation, Milford, MA) for reverse phase separations. The column was an in-house pulled-and-packed fused silica column with an I.D. of 100 microns pulled to a nanospray emitter.

Packing material for the column was C18 reprosil Pur 1.8 micron stationary phase (Dr. Maisch) with a total length of ∼25 cm. The UPLC system was set to a flow rate of 300 nL/min, where mobile phase A was 0.2% formic acid in water and mobile phase B was 0.2% formic acid in acetonitrile. Gel-extracted peptides were directly injected onto the column with a gradient of 2- 45% mobile phase B, followed by a high-B wash over a total 90 minutes. The mass spectrometer was operated in a data-dependent mode using HCD/ETD decision-tree fragmentation in the orbitrap (HCD) or ion trap (ETD) for MS/MS spectral generation.

The collected mass spectra were analyzed using Byonic v 2.12.0 or v2.14.27 (Protein Metrics) as the search engine for peptide ID and protein inference. The precursor and fragment ion tolerance were both set to 12 ppm and 0.4 Da for HCD and ETD, respectively. The search assumed tryptic digestion and allowed for up to two missed cleavages. Potential crosslinked peptides were identified using Byonic’s xlink feature, allowing for linker mass where appropriate. Data were validated using the standard reverse-decoy technique at a 1% false discovery rate, and inspected using Byologic for crosslink verification and assessment.

### Cysteine crosslinking

Symmetric inter-subunit ctSTIM1 dimers (5 μM) were incubated with 300 μM CuSO4 and 900 μM o-phenanthroline for 15 min at room temperature, after which the reaction was quenched with 2x LDS buffer containing 50 mM EDTA. After samples were run on an SDS-PAGE gel and stained with Coomassie Blue, gels were destained and scanned with a LI-COR Odyssey infrared imaging system. Crosslinking efficiency was quantified by the integrated intensity of the dimer band as a fraction of the summed intensities of the monomer and dimer bands.

For crosslinking flSTIM1 *in situ*, HEK293 cells were transfected with mCh-STIM1- C437S constructs with selected residues replaced by cysteine. 48 h after transfection, cells were exposed to 0.2 mM diamide for 10 min in either 2 Ca Ringer’s (full Ca^2+^ stores) or in 0 Ca^2+^ + 1 mM EGTA + 20 µM cyclopiazonic acid (depleted Ca^2+^ stores; CPA, Sigma). Cells were then lysed in RIPA containing 20 mM NEM, protease inhibitor cocktail (Cell Signaling Technology, #5871S) and EDTA. Samples were run on SDS-PAGE and analyzed by Western blot using mouse anti-mCherry primary antibody (1:2000, Takara Bio #632543) and Licor secondary antibody (Licor #926032212) on a LI-COR Odyssey imaging system. 50 mM DTT was added to duplicate samples to test for disulfide crosslinks. Sites of interest were replicated 2-3 times (biological replications with newly transfected cells).

### Calcium imaging

For experiments with STIM1 fragments (Supplementary Figure 7), HEK293 cells were transfected with mCherry-labeled STIM1 cytosolic fragments (0.25 µg + 0.25 µg empty pcDNA3 vector) and Orai1-GFP (0.5 µg) in a 35-mm dish using lipofectamine2000. Transfected cells were grown overnight in DMEM supplemented with Glutamax, 10% FBS, pen/strep, and 20 µM LaCl3 to minimize the toxic effects of constitutive Ca^2+^ influx. Cells were loaded with 1 µg/ml fura-2/AM for 30 min at room temperature, rinsed, and plated on poly-D-lysine-coated coverslips for imaging.

For experiments with flSTIM1, HEK293 cells were transfected with flSTIM1 cysteine mutants with Orai1-GFP. 24-48 h after transfection, cells were loaded with 1 µM fura-2/AM for 30 min at room temperature, rinsed, and plated on poly-D-lysine treated coverslip chambers.

Fura-2 imaging was conducted using a Zeiss 200M inverted microscope with a Fluar 40X NA 1.3 objective; cells were excited alternately at 350 and 380 nm (Polychrome II, TILL Photonics) and emission at 534 ± 30 nm (FF02-534/30-25, Semrock) or >480 nm (Chroma) was collected with a Flash4.0 sCMOS camera (Hamamatsu Corp.) with 2x2 binning. Images were background corrected before calculation of the 350/380 ratio for each individual cell. Standard bath solution contained (in mM): 155 NaCl, 4.5 KCl, 2 CaCl2, 1 MgCl2, 10 D-glucose and 5 Na-HEPES (pH 7.4). Ca^2+^-free solution was prepared by replacing CaCl2 with 2 mM MgCl2 and 1 mM EGTA. For cysteine crosslinking experiments, cells were treated with 1 µM ionomycin + 0.25 mM diamide in Ca^2+^-free Ringer’s followed by Ca^2+^-free Ringer’s + 0.25 mM diamide + 0.1% BSA to wash out ionomycin prior to readding standard (2 mM Ca^2+^) Ringer’s + 0.1% BSA. Calcium imaging was performed on a sufficient number of cells to obtain an error of the mean that allowed a clear distinction among responses to different conditions. Cells with an abnormally high or unstable baseline calcium level were excluded.

## Acknowledgements

The authors thank members of the Lewis lab for helpful discussions during the course of this project. This work was supported by NIH grant R37GM45374, the Mathers Charitable Foundation, and a Stanford Medicine Discovery Innovation Award (R.S.L.), and by NIH grant R37MH63105 (A.T.B.). S.v.D. was supported by Rubicon postdoctoral fellowship 825.13.027 from the Dutch Research Council NWO, and postdoctoral fellowship 16POST30780015 from the American Heart Association. Mass spectrometry was performed in collaboration with the Vincent Coates Foundation Mass Spectrometry Laboratory, Stanford University Mass Spectrometry. We thank R. Leib and K. Singhal for mass spectrometry analysis of crosslinked STIM1. The work was supported in part by NIH P30 CA124435 utilizing the Stanford Cancer Institute Proteomics/Mass Spectrometry Shared Resource.

## References

Bakan, A., L.M. Meireles, and I. Bahar. 2011. ProDy: Protein dynamics inferred from theory and experiments. Bioinformatics. 27:1575–1577. doi:10.1093/bioinformatics/btr168.

Böhm, J., and J. Laporte. 2018. Gain-of-function mutations in STIM1 and ORAI1 causing tubular aggregate myopathy and Stormorken syndrome. Cell Calcium. 76:1–9. doi:10.1016/j.ceca.2018.07.008.

Brunger, A.T. 2007. Version 1.2 of the Crystallography and NMR system. Nature protocols. 2:2728–33. doi:10.1038/nprot.2007.406.

Brunger, A.T., P.D. Adams, G.M. Clore, W.L. Delano, P. Gros, R.W. Grossekunstleve, J.S. Jiang, J. Kuszewski, M. Nilges, N.S. Pannu, R.J. Read, L.M. Rice, T. Simonson, and G.L. Warren. 1998. Crystallography & NMR system: A new software suite for macromolecular structure determination. Acta Crystallographica Section D: Biological Crystallography. 54:905–921. doi:10.1107/S0907444998003254.

Brunger, A.T., P. Strop, M. Vrljic, S. Chu, and K.R. Weninger. 2011. Three-dimensional molecular modeling with single molecule FRET. J Struct Biol. 173:497–505. doi:10.1016/j.jsb.2010.09.004.

Butorac, C., M. Muik, I. Derler, M. Stadlbauer, V. Lunz, A. Krizova, S. Lindinger, R. Schober, I. Frischauf, R. Bhardwaj, M. Hediger, K. Groschner, and C. Romanin. 2019. A novel STIM1-Orai1 gating interface essential for CRAC channel activation. Cell Calcium. 79:57– 67. doi:S0143416019300156.

Calloway, N.T., D.A. Holowka, and B.A. Baird. 2010. A basic sequence in STIM1 promotes Ca2+ influx by interacting with the C-terminal acidic coiled-coil of Orai1. Biochemistry. 49:1067–1071. doi:10.1021/bi901936q.

Choi, U.B., P. Strop, M. Vrljic, S. Chu, A.T. Brunger, and K.R. Weninger. 2010. Single- molecule FRET-derived model of the synaptotagmin 1-SNARE fusion complex. Nat Struct Mol Biol. 17:318–324. doi:10.1038/nsmb.1763.

Covington, E.D., M.M. Wu, and R.S. Lewis. 2010. Essential role for the CRAC activation domain in store-dependent oligomerization of STIM1. Mol Biol Cell. 21:1897–1907. doi:10.1091/mbc.E10-02-0145.

Cui, B., X. Yang, S. Li, Z. Lin, Z. Wang, C. Dong, and Y. Shen. 2013. The inhibitory helix controls the intramolecular conformational switching of the C-terminus of STIM1. PLoS One. 8:e74735. doi:10.1371/journal.pone.0074735.

Edelstein, A., N. Amodaj, K. Hoover, R. Vale, and N. Stuurman. 2010. Computer control of microscopes using µManager. Current protocols in molecular biology. 14:20.1-20.17. doi:10.1002/0471142727.mb1420s92.

Fahrner, M., M. Muik, R. Schindl, C. Butorac, P. Stathopulos, L. Zheng, I. Jardin, M. Ikura, and C. Romanin. 2014. A coiled-coil clamp controls both conformation and clustering of stromal interaction molecule 1 (STIM1). J Biol Chem. 289:33231–33244. doi:10.1074/jbc.M114.610022.

Fahrner, M., M. Stadlbauer, M. Muik, P. Rathner, P. Stathopulos, M. Ikura, N. Müller, and C. Romanin. 2018. A dual mechanism promotes switching of the Stormorken STIM1 R304W mutant into the activated state. Nat Commun. 9:825. doi:10.1038/s41467-018-03062-w.

Greenfeld, M., D.S. Pavlichin, H. Mabuchi, and D. Herschlag. 2012. Single Molecule Analysis Research Tool (SMART): an integrated approach for analyzing single molecule data. PLoS One. 7:e30024. doi:10.1371/journal.pone.0030024.

Grünberg, R., M. Nilges, and J. Leckner. 2006. Flexibility and Conformational Entropy in Protein-Protein Binding. Structure. 14:683–693. doi:10.1016/j.str.2006.01.014.

Harrison, S.C. 2008. Viral membrane fusion. Nature Structural and Molecular Biology. 15:690– 698. doi:10.1038/nsmb.1456.

Hirve, N., V. Rajanikanth, P.G. Hogan, and A. Gudlur. 2018. Coiled-coil formation conveys a STIM1 signal from ER lumen to cytoplasm. Cell Rep. 22:72–83. doi:10.1016/j.celrep.2017.12.030.

Kawasaki, T., I. Lange, and S. Feske. 2009. A minimal regulatory domain in the C terminus of STIM1 binds to and activates ORAI1 CRAC channels. Biochem Biophys Res Commun. 385:49–54. doi:10.1016/j.bbrc.2009.05.020.

Komolov, K.E., Y. Du, N.M. Duc, R.M. Betz, J.P.G.L.M. Rodrigues, R.D. Leib, D. Patra, G. Skiniotis, C.M. Adams, R.O. Dror, K.Y. Chung, B.K. Kobilka, and J.L. Benovic. 2017. Structural and Functional Analysis of a β2-Adrenergic Receptor Complex with GRK5. Cell. 169:407–421.e16. doi:10.1016/j.cell.2017.03.047.

Korzeniowski, M.K., I.M. Manjarrés, P. Varnai, and T. Balla. 2010. Activation of STIM1-Orai1 involves an intramolecular switching mechanism. Sci Signal. 3:ra82. doi:10.1126/scisignal.2001122.

Lacruz, R.S., and S. Feske. 2015. Diseases caused by mutations in ORAI1 and STIM1. Ann NY Acad Sci. 1356:45–79. doi:10.1111/nyas.12938.

Liou, J., M. Fivaz, T. Inoue, and T. Meyer. 2007. Live-cell imaging reveals sequential oligomerization and local plasma membrane targeting of stromal interaction molecule 1 after Ca^2+^ store depletion. Proc Natl Acad Sci USA. 104:9301–9306.

London, N., B. Raveh, E. Cohen, G. Fathi, and O. Schueler-Furman. 2011. Rosetta FlexPepDock web server - high resolution modeling of peptide-protein interactions. Nucleic Acids Research. 39:249–253. doi:10.1093/nar/gkr431.

Ma, G., M. Wei, L. He, C. Liu, B. Wu, S.L. Zhang, J. Jing, X. Liang, A. Senes, P. Tan, S. Li, A. Sun, Y. Bi, L. Zhong, H. Si, Y. Shen, M. Li, M.-S. Lee, W. Zhou, J. Wang, Y. Wang, and Y. Zhou. 2015. Inside-out Ca^2+^ signalling prompted by STIM1 conformational switch. Nat Commun. 6:7826. doi:10.1038/ncomms8826.

McCann, J.J., U.B. Choi, L. Zheng, K. Weninger, and M.E. Bowen. 2010. Optimizing methods to recover absolute FRET efficiency from immobilized single molecules. Biophys J. 99:961–970. doi:10.1016/j.bpj.2010.04.063.

Misceo, D., A. Holmgren, W.E. Louch, P.A. Holme, M. Mizobuchi, R.J. Morales, A.M. De Paula, A. Stray-Pedersen, R. Lyle, B. Dalhus, G. Christensen, H. Stormorken, G.E. Tjønnfjord, and E. Frengen. 2014. A Dominant STIM1 Mutation Causes Stormorken Syndrome. Human Mutation. 35:556–564. doi:10.1002/humu.22544.

Morin, G., N.O. Bruechle, A.R. Singh, C. Knopp, G. Jedraszak, M. Elbracht, D. Brémond- Gignac, K. Hartmann, H. Sevestre, P. Deutz, D. Hérent, P. Nürnberg, B. Roméo, K. Konrad, M. Mathieu-Dramard, J. Oldenburg, E. Bourges-Petit, Y. Shen, K. Zerres, H. Ouadid-Ahidouch, and J. Rochette. 2014. Gain-of-Function Mutation in STIM1 (P.R304W) Is Associated with Stormorken Syndrome. Hum Mutat. 35:1221–1232. doi:10.1002/humu.22621.

Muik, M., M. Fahrner, I. Derler, R. Schindl, J. Bergsmann, I. Frischauf, K. Groschner, and C. Romanin. 2009. A cytosolic homomerization and a modulatory domain within STIM1 C terminus determine coupling to ORAI1 channels. J Biol Chem. 284:8421–8426. doi:10.1074/jbc.C800229200.

Muik, M., M. Fahrner, R. Schindl, P. Stathopulos, I. Frischauf, I. Derler, P. Plenk, B. Lackner, K. Groschner, M. Ikura, and C. Romanin. 2011. STIM1 couples to ORAI1 via an intramolecular transition into an extended conformation. EMBO J. 30:1678–1689. doi:10.1038/emboj.2011.79.

Nesin, V., G. Wiley, M. Kousi, E.-C. Ong, T. Lehmann, D.J. Nicholl, M. Suri, N. Shahrizaila, N. Katsanis, P.M. Gaffney, K.J. Wierenga, and L. Tsiokas. 2014. Activating mutations in STIM1 and ORAI1 cause overlapping syndromes of tubular myopathy and congenital miosis. Proc Natl Acad Sci USA. 111:4197–4202. doi:10.1073/pnas.1312520111.

Park, C.Y., P.J. Hoover, F.M. Mullins, P. Bachhawat, E.D. Covington, S. Raunser, T. Walz, K.C. Garcia, R.E. Dolmetsch, and R.S. Lewis. 2009. STIM1 clusters and activates CRAC channels via direct binding of a cytosolic domain to Orai1. Cell. 136:876–890. doi:10.1016/j.cell.2009.02.014.

Prakriya, M., and R.S. Lewis. 2015. Store-operated calcium channels. Physiol Rev. 95:1383– 1436. doi:10.1152/physrev.00020.2014.

Qin, M., W. Wang, and D. Thirumalai. 2015. Protein folding guides disulfide bond formation. Proc Natl Acad Sci USA. 112:11241–11246. doi:10.1073/pnas.1503909112.

Rappsilber, J. 2011. The beginning of a beautiful friendship: cross-linking/mass spectrometry and modelling of proteins and multi-protein complexes. J Struct Biol. 173:530–540. doi:10.1016/j.jsb.2010.10.014.

Rathner, P., M. Fahrner, L. Cerofolini, H. Grabmayr, F. Horvath, H. Krobath, A. Gupta, E. Ravera, M. Fragai, M. Bechmann, T. Renger, C. Luchinat, C. Romanin, and N. Müller. 2020. Interhelical interactions within the STIM1 CC1 domain modulate CRAC channel activation. Nature Chemical Biology. doi:10.1038/s41589-020-00672-8.

Roy, R., S. Hohng, and T. Ha. 2008. A practical guide to single-molecule FRET. Nat Methods. 5:507–516. doi:10.1038/nmeth.1208.

Schmidt, C., and C. V Robinson. 2014. A comparative cross-linking strategy to probe conformational changes in protein complexes. Nat Protoc. 9:2224–2236. doi:10.1038/nprot.2014.144.

Schober, R., D. Bonhenry, V. Lunz, J. Zhu, A. Krizova, I. Frischauf, M. Fahrner, M.Q. Zhang, L. Waldherr, T. Schmidt, I. Derler, P.B. Stathopulos, C. Romanin, R.H. Ettrich, and R. Schindl. 2019. Sequential activation of STIM1 links Ca^2+^ with luminal domain unfolding. Sci Signal. 12. doi:10.1126/scisignal.aax3194.

Soboloff, J., B.S. Rothberg, M. Madesh, and D.L. Gill. 2012. STIM proteins: dynamic calcium signal transducers. Nat Rev Mol Cell Biol. 13:549–565. doi:10.1038/nrm3414.

Stathopulos, P., G. Li, M. Plevin, J. Ames, and M. Ikura. 2006. Stored Ca^2+^ depletion-induced oligomerization of STIM1 via the EF-SAM region: An initiation mechanism for capacitive Ca^2+^ entry. J Biol Chem. 281:35855–35862.

Stathopulos, P.B., R. Schindl, M. Fahrner, L. Zheng, G.M. Gasmi-Seabrook, M. Muik, C. Romanin, and M. Ikura. 2013. STIM1/Orai1 coiled-coil interplay in the regulation of store- operated calcium entry. Nat Commun. 4:2963. doi:10.1038/ncomms3963.

Stryer, L., and R.P. Haugland. 1967. Energy transfer: a apectroscopic ruler. 58:719–726.

Thompson, J.L., Y. Zhao, P.B. Stathopulos, A. Grossfield, and T.J. Shuttleworth. 2018. Phosphorylation-mediated structural changes within the SOAR domain of STIM1 enable specific activation of distinct Orai channels. J Biol Chem. 293:3145–3155. doi:10.1074/jbc.M117.819078.

Wang, X., Y. Wang, Y. Zhou, E. Hendron, S. Mancarella, M.D. Andrake, B.S. Rothberg, J. Soboloff, and D.L. Gill. 2014. Distinct Orai-coupling domains in STIM1 and STIM2 define the Orai-activating site. Nat Commun. 5:3183. doi:10.1038/ncomms4183.

Weiss, S. 1999. Fluorescence spectroscopy of single biomolecules. Science. 283:1676–1683. doi:10.1126/science.283.5408.1676.

Weiss, S. 2000. Measuring conformational dynamics of biomolecules by single molecule fluorescence spectroscopy. Nature Structural Biology. 7:724–729. doi:10.1038/78941.

Wu, M.M., J. Buchanan, R.M. Luik, and R.S. Lewis. 2006. Ca^2+^ store depletion causes STIM1 to accumulate in ER regions closely associated with the plasma membrane. J Cell Biol. 174:803–813. doi:10.1083/jcb.200604014.

Wu, M.M., E.D. Covington, and R.S. Lewis. 2014. Single-molecule analysis of diffusion and trapping of STIM1 and Orai1 at endoplasmic reticulum-plasma membrane junctions. Mol Biol Cell. 25:3672–3685. doi:10.1091/mbc.E14-06-1107.

Yang, X., H. Jin, X. Cai, S. Li, and Y. Shen. 2012. Structural and mechanistic insights into the activation of Stromal interaction molecule 1 (STIM1). Proc Natl Acad Sci USA. 109:5657– 5662. doi:10.1073/pnas.1118947109.

Yu, F., L. Sun, S. Hubrack, S. Selvaraj, and K. Machaca. 2013. Intramolecular shielding maintains the ER Ca^2+^ sensor STIM1 in an inactive conformation. J Cell Sci. 126:2401– 2410. doi:10.1242/jcs.117200.

Yuan, J., W. Zeng, M. Dorwart, Y. Choi, P. Worley, and S. Muallem. 2009. SOAR and the polybasic STIM1 domains gate and regulate Orai channels. Nat Cell Biol. 11:337–343. doi:10.1038/ncb1842.

Zhou, Y., P. Srinivasan, S. Razavi, S. Seymour, P. Meraner, A. Gudlur, P.B. Stathopulos, M. Ikura, A. Rao, and P.G. Hogan. 2013. Initial activation of STIM1, the regulator of store- operated calcium entry. Nat Struct Mol Biol. 20:973–981. doi:10.1038/nsmb.2625.

