## Supplementary figures for "Conformational dynamics of auto-inhibition in the ER calcium sensor STIM1"

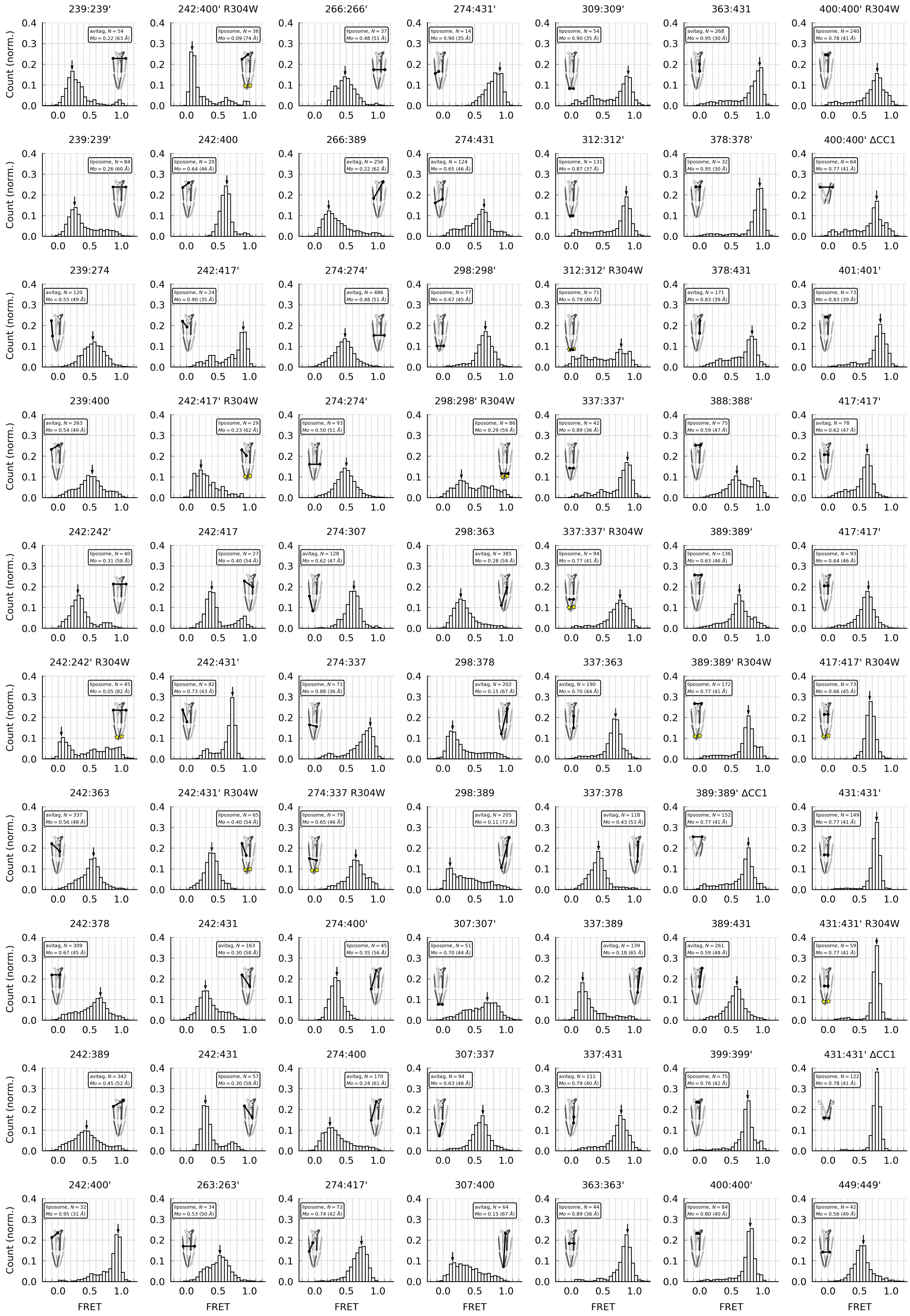


**Supplementary Figure 1. smFRET histograms.** All smFRET measurements used in this study. Peaks of the histograms are indicated by an arrow and were considered to represent the predominant ctSTIM1 conformation. Homomeric and heteromeric ctSTIM1 cysteine mutants were designed to enable symmetric inter-subunit smFRET, asymmetric inter-subunit smFRET, and intra-subunit smFRET. The locations of the measurement sites are indicated on a model of ctSTIM1 or CAD for each sample. Legend boxes indicate immobilization method (liposome vs avitag), number of molecules (N) used to construct the histogram, the peak FRET value (Mo) and the corresponding distance (using $R_0$=5.1 nm). Liposome encapsulation and direct surface attachment gave similar results for inter-subunit smFRET at 239:239′, 274:274′, and 417:417′, and for intra-subunit smFRET at 242:431. CC1:CC1′ and CC1:CAD measurements (36 unique site combinations) were used to constrain the CC1-CAD model. For sites with both liposome and avitag measurements, the liposome measurement was used.

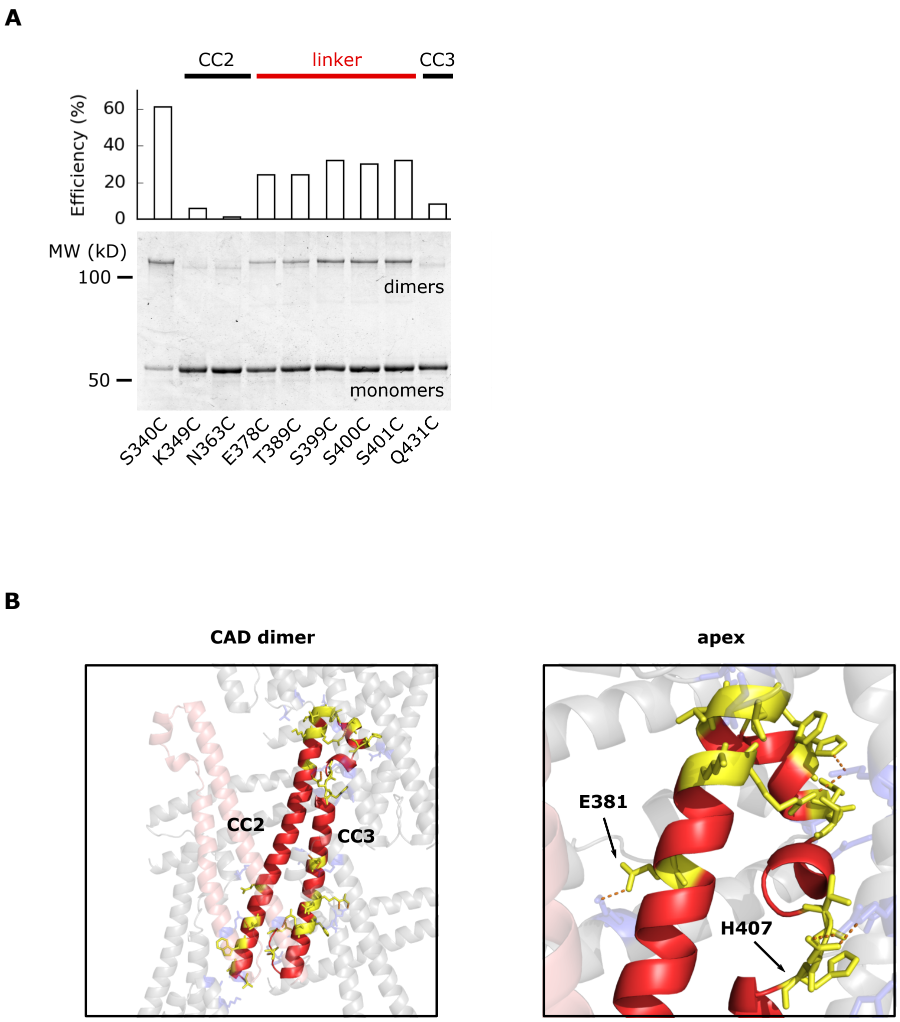


**Supplementary Figure 2. CAD apex structure.** (**A**) Dimerization of ctSTIM1 cysteine mutants by disulfide bond formation in CAD, shown by non-reducing SDS-PAGE. CuP-induced dimerization was virtually absent at basal sites but was evident at all tested sites in the apical region. (**B**) Analysis of external contacts of a CAD subunit in the reconstructed crystal lattice. In the left panel a subunit of CAD is shown in red, the adjacent subunit in the dimer is shown in transparent pink, and interacting subunits in the crystal lattice are shown in transparent gray. The crystal lattice was generated from structural coordinates in 3TEQ.pdb using Chimera software (Pettersen et al., 2004). Residues on the red subunit that are in contact with the external crystal lattice (including the paired subunit) are shown in yellow, and their corresponding contacts in the external lattice are shown in blue. Contacts are defined as sites where Van der Waals radii touch or overlap. Extensive external contacts are made at the base of CAD, near the dimer interface, and at the apex in the CC2-CC3 linker region. The enlargement on the right shows how lattice contacts are made virtually throughout the entire apex region. Polar contacts (hydrogen bonds) are shown as dashed orange lines.

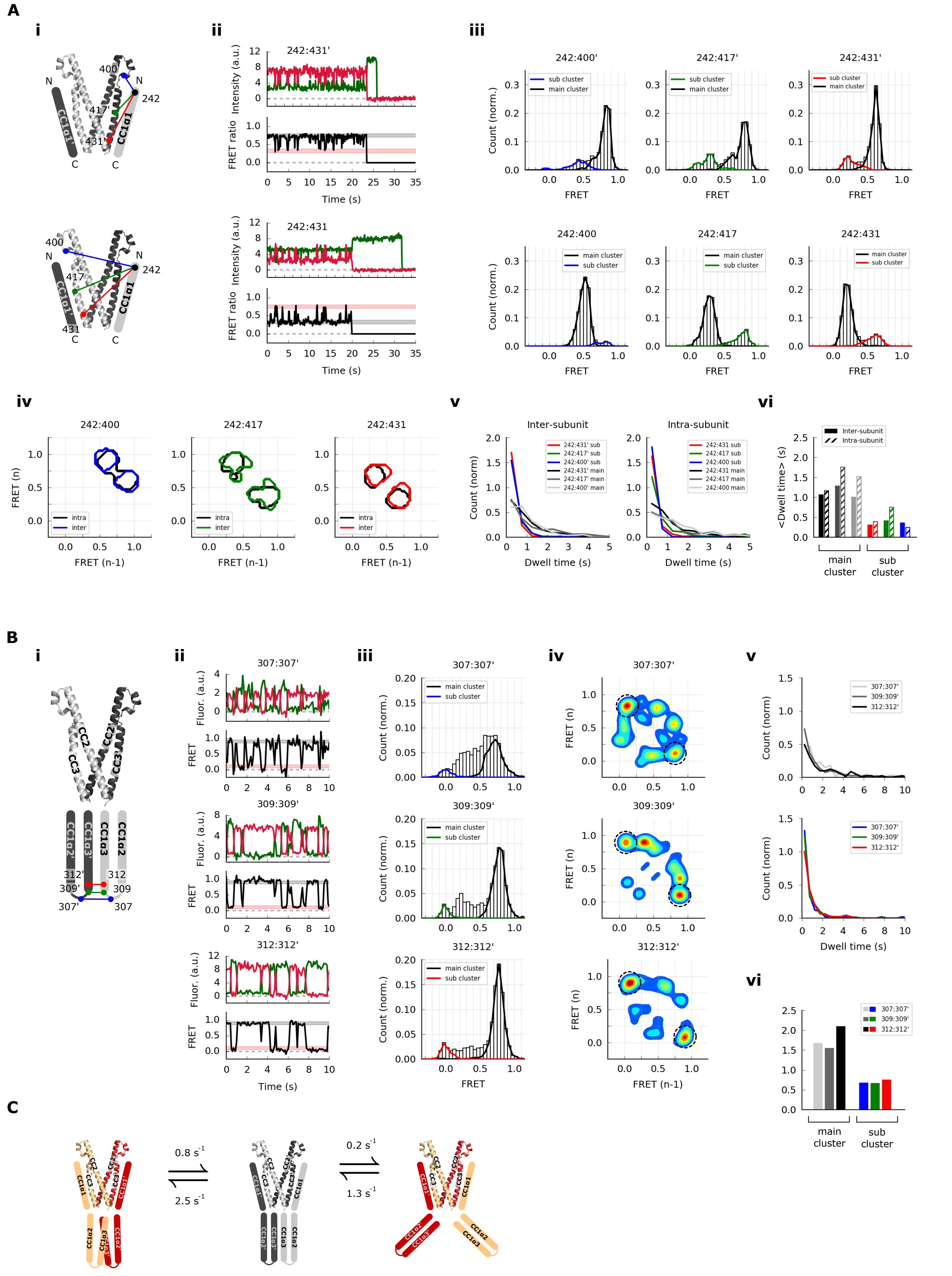


**Supplementary Figure 3. CC1 FRET fluctuations.** (**A**) CC1α1:CAD′ fluctuations. (**Ai**) Schematic drawing of the measurement sites, comparing inter-subunit (*top*) and intra-subunit (*bottom*) smFRET measurements from aa 242 in CC1α1 to sites 400, 417, and 431 in CC3. (**Aii**) Examples of smFRET transitions at 242:431′ (inter-subunit, *top*) and 242:431 (intra-subunit, *bottom*). Black and red bars indicate the predominant and transient FRET levels, respectively. (**Aiii**) Ensemble smFRET histograms show the equivalence of predominant and transient FRET levels in inter- and intra-subunit measurements, suggesting that CC1α1 transiently switched sides on CAD, probably using the same binding interface. (**Aiv**) Ensemble transition density plots, plotting FRET level after a transition against FRET before a transition, for all fluctuating molecules (see Methods). Contours indicate which transitions were most often observed, overlaying intra-subunit (*black*) and inter-subunit (*colored*) measurements. (**Av**) Dwell time distributions for inter-subunit (*left*) and intra-subunit (*right*) measurements were similar, further supporting the hypothesis that CC1α1 domains transiently switched sides on CAD. (**Avi**) Average dwell times were similar between the various measurement configurations, for predominant and transient FRET levels. Average dwell times of the predominant and transient states were 1.2 s and 0.4 s, respectively. (**B**) CC1α3:CC1α3′ fluctuations. (**Bi**) Schematic drawing of measurement sites, comparing inter-subunit smFRET measurements at sites near the CC1α3 N-terminus. (**Bii**) Example traces at aa 307 (*top*), 309 (*center*), and 312 (*bottom*) show large-amplitude fluctuations from the predominant high FRET level. Black and red bars indicate the predominant and transient FRET levels, respectively. (**Biii**) Ensemble smFRET histograms with predominant and transient FRET clusters associated with the large-amplitude fluctuations. (**Biv**) FRET transition diagrams show that predominant fluctuations at all three sites are large-amplitude transitions between a high- and a low-FRET state (*dashed circles*). (**Bv**) Dwell time distributions of the predominant FRET state (*top*) and the transient state (*bottom*) were similar for all three sites, indicating a common underlying conformational change. (**Bvi**) Average dwell times for all three sites. Overall average dwell times were 1.7 s for the predominant state, and 0.7 s for the transient sub-state. (**C**) Two types of conformational changes in the CC1 domain. From the predominant resting state (*center*), CC1α1 domains occasionally switch sides on CAD (*left*), and CC1α3 domains occasionally open up (*right*). As their kinetic rates appear different, the two conformational transitions are apparently not directly coupled, and the CC1α2/3 domains may maintain a compact folding during the CC1α1 transition.

**A**

| | FRET pair | FRET peak | Dist. peak (Å) | Dist. low (Å) | Dist. high (Å) | Δ low (Å) | Δ high (Å) |
|---|---|---|---|---|---|---|---|
| 1 | **239:239'** | 0.26 | 60 | 57 | 64 | -20 | 0 |
| 2 | **239:274** | 0.55 | 49 | 47 | 51 | -20 | 0 |
| 3 | **239:400** | 0.54 | 49 | 47 | 51 | -10 | 0 |
| 4 | **242:242'** | 0.31 | 58 | 55 | 61 | -20 | 0 |
| 5 | **242:363** | 0.56 | 48 | 46 | 50 | -10 | 0 |
| 6 | **242:378** | 0.67 | 45 | 43 | 47 | -10 | 0 |
| 7 | **242:389** | 0.45 | 52 | 50 | 54 | -10 | 0 |
| 8 | **242:400** | 0.64 | 46 | 44 | 48 | -10 | 0 |
| 9 | **242:400'** | 0.95 | 31 | 0 | 36 | 0 | 0 |
| 10 | **242:417** | 0.40 | 54 | 52 | 56 | -10 | 0 |
| 11 | **242:417'** | 0.90 | 35 | 30 | 38 | -10 | 0 |
| 12 | **242:431** | 0.30 | 58 | 56 | 61 | -10 | 0 |
| 13 | **242:431'** | 0.73 | 43 | 40 | 45 | -10 | 0 |
| 14 | **266:266'** | 0.48 | 51 | 49 | 53 | -20 | 0 |
| 15 | **266:389** | 0.22 | 62 | 59 | 67 | -10 | 0 |
| 16 | **274:274'** | 0.50 | 51 | 48 | 53 | -20 | 0 |
| 17 | **274:307** | 0.62 | 47 | 45 | 49 | -20 | 0 |
| 18 | **274:337** | 0.88 | 36 | 32 | 39 | -20 | 0 |
| 19 | **274:400** | 0.24 | 61 | 58 | 65 | -10 | 0 |
| 20 | **274:400'** | 0.35 | 56 | 54 | 59 | -10 | +10 |
| 21 | **274:417'** | 0.74 | 42 | 40 | 44 | -10 | 0 |
| 22 | **274:431** | 0.65 | 46 | 44 | 48 | -10 | 0 |
| 23 | **274:431'** | 0.90 | 35 | 30 | 38 | -10 | 0 |
| 24 | **298:298'** | 0.67 | 45 | 43 | 47 | -20 | 0 |
| 25 | **298:363** | 0.28 | 59 | 56 | 62 | -10 | +10 |
| 26 | **298:378** | 0.15 | 67 | 63 | 74 | -10 | +20 |
| 27 | **298:389** | 0.11 | 72 | 66 | 84 | -10 | +30 |
| 28 | **307:337** | 0.63 | 46 | 44 | 48 | -20 | 0 |
| 29 | **307:400** | 0.15 | 67 | 63 | 74 | -10 | +40 |
| 30 | **309:309'** | 0.90 | 35 | 30 | 38 | -20 | 0 |
| 31 | **312:312'** | 0.87 | 37 | 33 | 40 | -20 | 0 |
| 32 | **337:337'** | 0.89 | 36 | 31 | 39 | -20 | 0 |
| 33 | **337:363** | 0.70 | 44 | 42 | 46 | -10 | 0 |
| 34 | **337:378** | 0.43 | 53 | 51 | 55 | -10 | +10 |
| 35 | **337:389** | 0.18 | 65 | 61 | 70 | -10 | +20 |
| 36 | **337:431** | 0.79 | 40 | 38 | 43 | -10 | 0 |

**B**

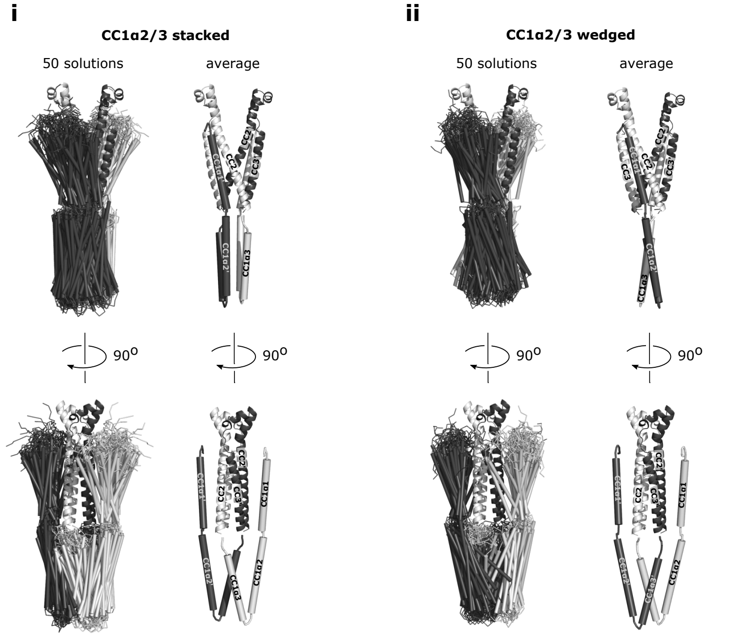


**C**

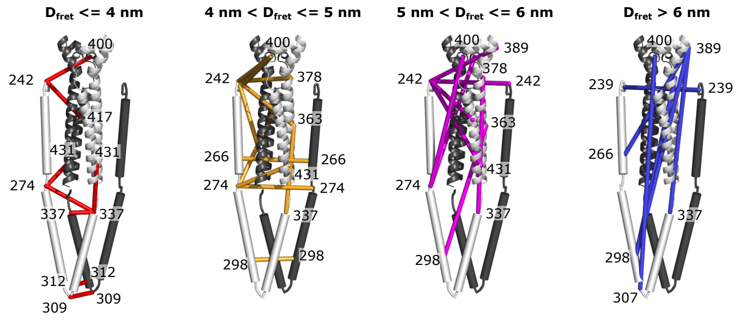


**Supplementary Figure 4. CC1-CAD model.** (**A**) Table of CC1:CC1 and CC1:CAD smFRET measurements used to construct the CC1-CAD model. The 'FRET peak' and 'Dist. peak' columns correspond to the peaks of the smFRET histograms, and their associated calculated distances, shown in Supplementary Figure 1. The 'Dist. low' and 'Dist. high' columns are lower and upper bounds of the distance range, based on an uncertainty of ±0.05 in the smFRET measurement. The 'Δ low' and 'Δ high' columns are adjustments to the distance bounds, based on a ~1 nm fluorophore linker length, to account for the absence of explicit fluorophore models in the CC1 domain (see Methods). Additional relaxation of the upper distance bound was allowed for some low-FRET measurements. The reason is that zero-FRET traces were excluded from the FRET histograms, resulting in over-estimated peak FRET levels (under-estimated distances) for such low-FRET measurement sites. This can have an especially large effect when the true FRET is very low (<0.1), as the smFRET technique is not sensitive in this range. (**B**) Results of the smFRET-constrained modeling. CC1α1 domains were oriented parallel to CC3 on the adjacent subunit. CC1α2/3 domains were compact, parallel-folded, and oriented away from CAD. The modeling produced two different topologies for stacking of the CC1α2/3 domains. (**Bi**) Overlay (*left*) and average (*right*) of 50 solutions with stacked CC1α2/3 domains. This was the predominant topology, representing ~70% of solutions. (**Bii**) Overlay (*left*) and average (*right*) of 50 solutions for the wedged CC1α2/3 topology. The orientation of CC1α1 domains was virtually identical for both classes of solutions. (**C**) FRET measurement sites indicated on the smFRET-derived model, grouped by their corresponding smFRET-derived distances $D_{FRET}$.

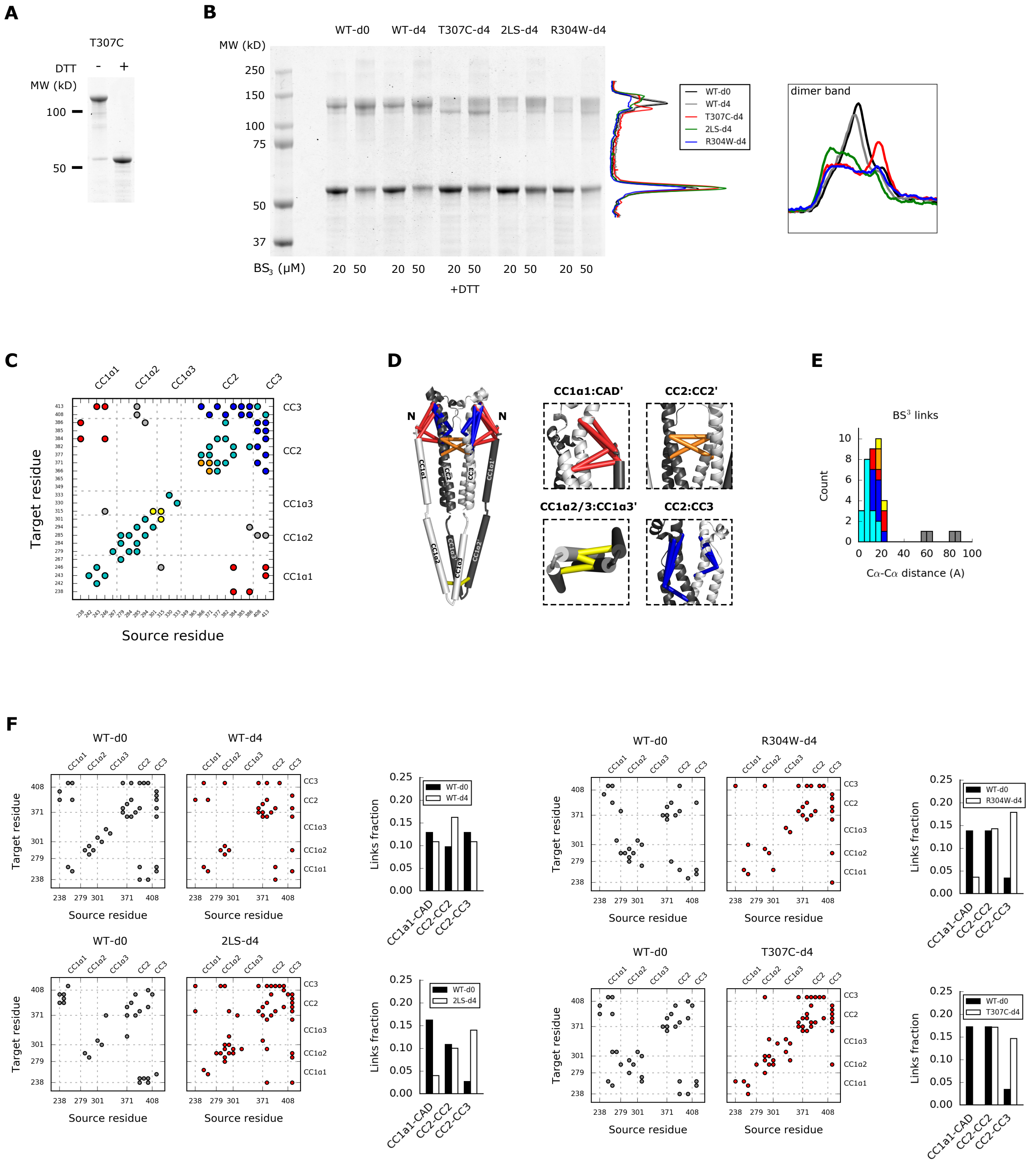


**Supplementary Figure 5. Mass spectrometry analysis of BS³ crosslinking. (A)** SDS-PAGE gel analysis of cysteine crosslinking at T307C using copper/phenanthroline (CuP). Crosslinking of T307C could be driven to almost 100% by one-hour exposure to CuP and could be completely reversed by addition of reducing agent DTT. (**B**) SDS-PAGE gel analysis of ctSTIM1 variants treated with BS³ for lysine-lysine crosslinking. Wild-type ctSTIM1 was treated with BS³-d0 to serve as a reference, and four different ctSTIM1 variants were treated with BS³-d4 to detect the effect of the variation using comparative XLMS (see Methods). The variants are a wild-type ctSTIM1 as control (WT-d4), a T307C cysteine-cysteine crosslinked ctSTIM1 (T307C-d4), and ctSTIM1 with activating mutations L248S+L251S (2LS-d4) and R304W (R304W-d4). The T307C sample was first cysteine-crosslinked with CuP (see **A**), before treatment with $BS^3$-d4. After $BS^3$-d4 treatment the cysteine crosslink was reversed by treatment with DTT. $BS^3$ treatment resulted in the formation of ctSTIM1 dimers, with distinct gel band patterns for wild-types and mutants. The integrated gel lane signals are shown on the right. (**C**) Identification of $BS^3$-linked lysine residues in CC1-CAD, detected by mass spectrometry analysis of wild-type ctSTIM1 monomer and dimer gel bands, cut from the SDS page gel. Identified links are indicated in a matrix of all possible lysine pairs in CC1-CAD. Links found exclusively in the dimer band were likely inter-subunit links (*red*, *orange*, or *yellow*). Links found in the monomer band were intra-subunit links (*blue* or *cyan*). (**D**) Mapping of detected $BS^3$ links on the smFRET-derived structural model. Inter-subunit links were detected for CC1α1:CAD′ (*red*), CC2:CC2′ (*orange*), and CC1α2:CC1α3′ (*yellow*). Intra-subunit links were detected for CC2:CC3 (*blue*), and for several sites that were in close proximity on the peptide chain (*cyan*). (**E**) Distribution of distances between residues for all detected links mapped on the smFRET-derived model (using the Cα atom for residues in CAD). The majority of linked residues matched well with the theoretical $BS^3$ length of ~15 Å, except links shown in gray. (**F**) Mass-spectrometry analysis of three ctSTIM1 variants (T307C, 2LS, and R304W) showed similarly altered link patterns. Relative to the WT reference, the number of CC1α1-CAD links decreased in the variants, consistent with the release of CC1α1 in an activated state of ctSTIM1. The decrease in CC1α1-CAD links was compensated by an increase in CC2-CC3 links within CAD, probably because the CC1α1 domain was no longer present to compete for links with sites in CAD.

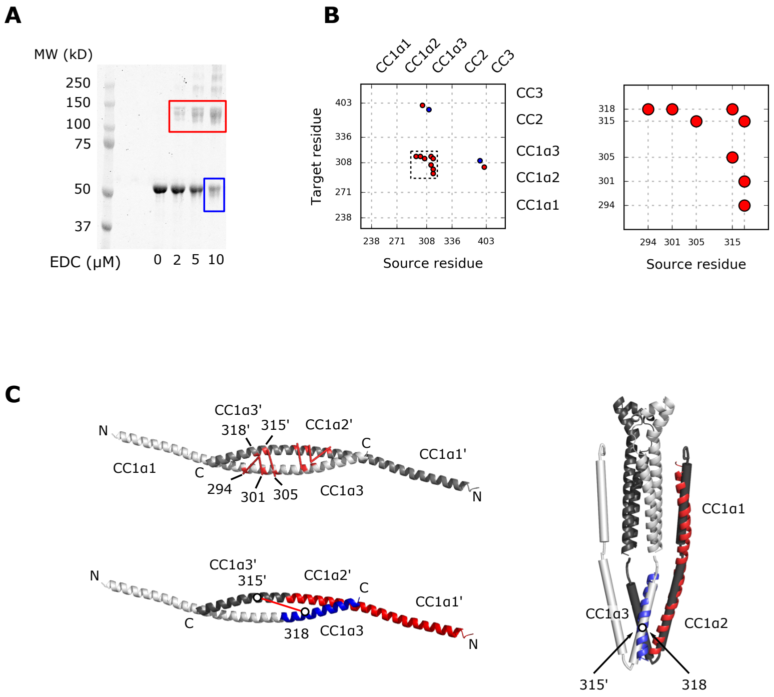


**Supplementary Figure 6. Mass spectrometry analysis of EDC crosslinking.** (**A**) SDS-PAGE gel analysis of ctSTIM1 treated with zero-length amine-carboxyl crosslinker EDC (see Methods). Monomer (*blue*) and dimer (*red*) bands were cut from the gel for analysis by mass spectrometry. For the monomer band we used only the last lane, to have approximately equal amounts of material from both bands and increase the fraction of reacted sample. (**B**) Mass spectrometry analysis showed a cluster of four dimeric (likely inter-subunit) links between CC1α2 and CC1α3 domains, indicated by the black dashed square and enlarged on the right. (**C**) A crystal structure of isolated CC1 fragment (237-340, with mutations M244L and L321M) is a dimer of two continuous helices with a coiled-coil configuration involving CC1α2 and CC1α3 domains (Cui et al., 2013). Three out of four EDC links in the CC1α2/3 cluster (K294:E318′, K301:E318′, and E305:K315′) could be mapped directly onto the crystal structure (*top left*). The fourth link (K315′:E318) appears to bridge a larger distance (*bottom left*), but could be rationalized by aligning the CC1 dimer crystal to the smFRET-derived CC1-CAD model and folding the CC1 dimer around the flexible region aa 305-310 (*right*). In the aligned CC1 dimer, residues 315′ and 318 are right next to each other.

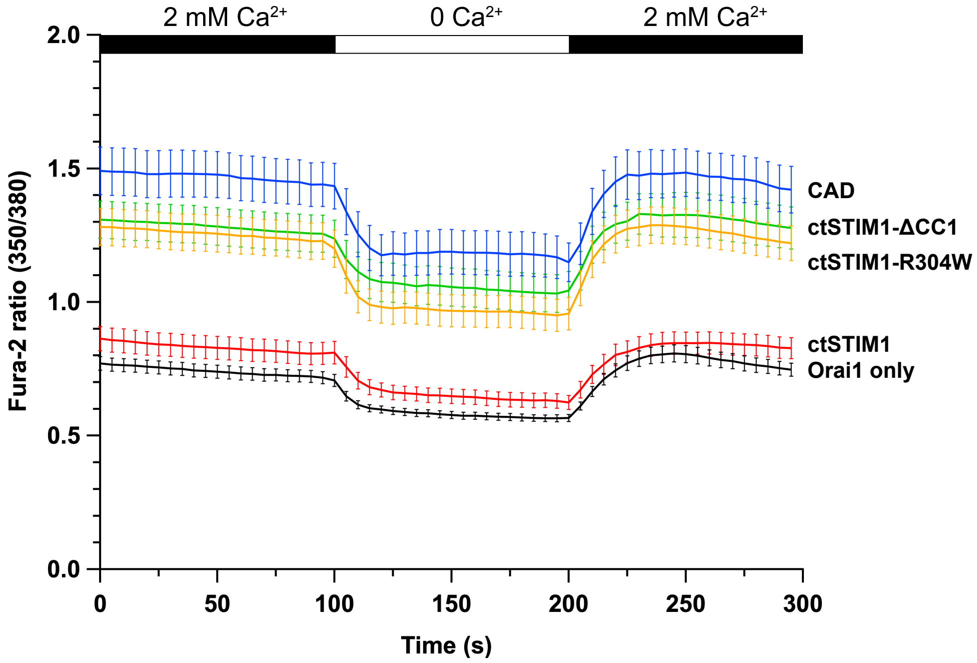


**Supplementary Figure 7. Comparison of SOCE evoked by ctSTIM1 fragments in HEK293 cells.** Cells expressing the indicated ctSTIM1 variant and Orai1-GFP were exposed sequentially to 2 mM $Ca^{2+}$, 0 $Ca^{2+}$, and 2 mM $Ca^{2+}$. The 350/380 fluorescence ratio is shown for ctSTIM1 (aa 233-685), ctSTIM1-R304W, ctSTIM1-ΔCC1 (aa 340-685), CAD (aa 342-448), and Orai1-GFP alone. Each trace shows the mean and s.e.m. of 25-35 cells in multiple runs from two independent transfections. ctSTIM1 shows very little activity, while ctSTIM1-R304W and ctSTIM1-ΔCC1 approach the high activity of the CAD fragment.

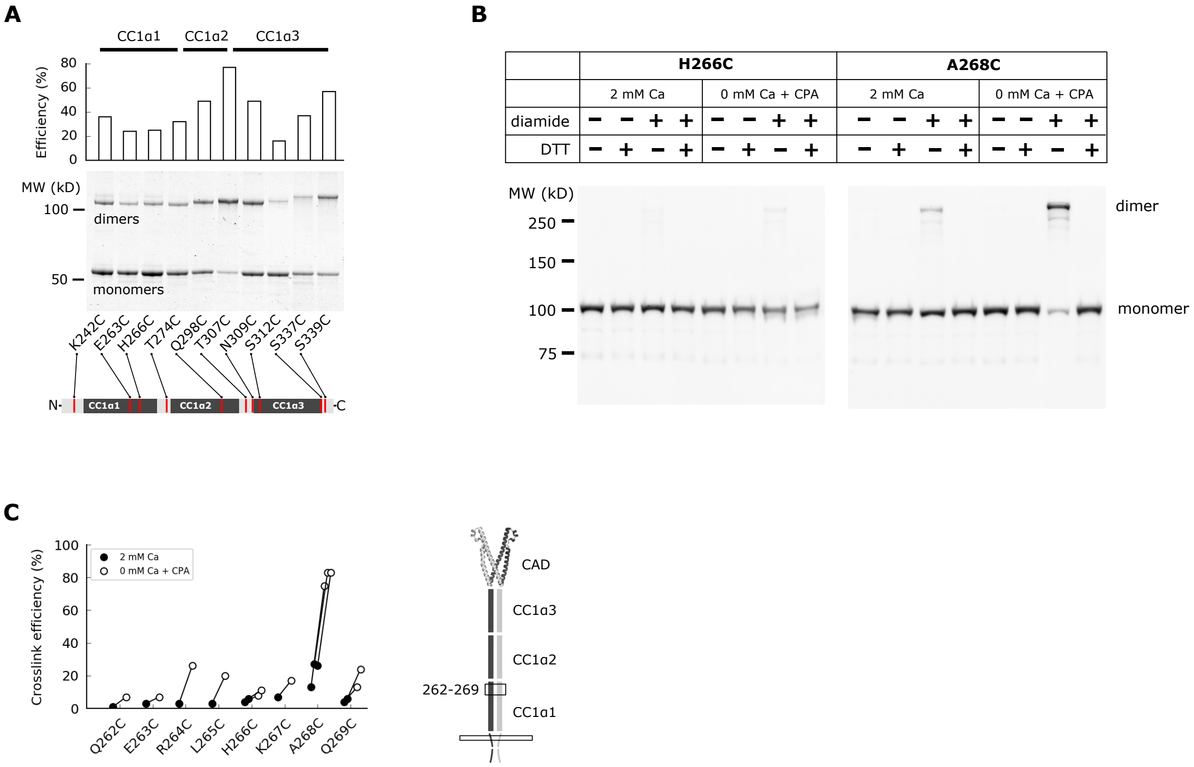


**Supplementary Figure 8. CC1 cysteine crosslinking in ctSTIM1 and flSTIM1.** (**A**) Cysteine crosslinking throughout the CC1 domain of ctSTIM1 mutants *in vitro* detected by non-reducing SDS-PAGE. Crosslinking efficiency peaked at aa 307 in the CC1α2/3 linker region. **(B)** Full-length STIM1 (flSTIM1) mutants, each containing a single introduced cysteine, were transiently over-expressed in HEK293 cells for diamide-induced cysteine crosslinking *in vivo*. Western-blot analysis of cysteine crosslinking of flSTIM1-H266C (*left*) and flSTIM1-A268C (*right*) from cells incubated with diamide, either under resting (2 mM Ca) or store-depleted (0 mM Ca + CPA) conditions. flSTIM1-A268C had very strong crosslinking in the activated state. (**C**) Crosslinking efficiencies in the inactive (*black*) and active (*white*) states for residues aa 262-269 at the C-terminal end of CC1α1. Crosslinking at sites within this region increased somewhat in the active state, but crosslinking at aa 268 was particularly strong, suggesting a direct, stable apposition at that position.

### References


Cui, B., X. Yang, S. Li, Z. Lin, Z. Wang, C. Dong, and Y. Shen. 2013. The inhibitory helix controls the intramolecular conformational switching of the C-terminus of STIM1. *PLoS One*. 8:e74735. doi:10.1371/journal.pone.0074735.

Pettersen, E.F., T.D. Goddard, C.C. Huang, G.S. Couch, D.M. Greenblatt, E.C. Meng, and T.E. Ferrin. 2004. UCSF Chimera - A visualization system for exploratory research and analysis. *Journal of Computational Chemistry*. 25:1605–1612. doi:10.1002/jcc.20084.
